# Immunofocusing on the conserved fusion peptide of HIV envelope glycoprotein in rhesus macaques

**DOI:** 10.1101/2024.11.27.625755

**Authors:** Payal P. Pratap, Christopher A. Cottrell, James Quinn, Diane G. Carnathan, Daniel L.V. Bader, Andy S. Tran, Chiamaka A. Enemuo, Julia T. Ngo, Sara T. Richey, Hongmei Gao, Xiaoying Shen, Kelli M. Greene, Jonathan Hurtado, Katarzyna Kaczmarek Michaels, Elana Ben-Akiva, Ashley Lemnios, Mariane B. Melo, Joel D. Allen, Gabriel Ozorowski, Max Crispin, Bryan Briney, David Montefiori, Guido Silvestri, Darrell J. Irvine, Shane Crotty, Andrew B. Ward

## Abstract

**Summary:** During infection, the fusion peptide (FP) of HIV envelope glycoprotein (Env) serves a central role in viral fusion with the host cell. As such, the FP is highly conserved and therefore an attractive epitope for vaccine design. Here, we describe a vaccination study in non-human primates (NHPs) where glycan deletions were made on soluble HIV Env to increase FP epitope exposure. When delivered via implantable osmotic pumps, this immunogen primed immune responses against the FP, which were then boosted with heterologous trimers resulting in a focused immune response targeting the conserved FP epitope. Although autologous immunizations did not elicit high affinity FP-targeting antibodies, the conserved FP epitope on a heterologous trimer further matured the lower affinity, FP-targeting B cells. This study suggests using epitope conservation strategies on distinct Env trimer immunogens can focus humoral responses on desired neutralizing epitopes and suppress immune-distracting antibody responses against non-neutralizing epitopes.

## Introduction

Broadly neutralizing antibodies (bnAbs) against HIV-1 envelope glycoprotein (Env) have been isolated from patients infected with HIV, but only after prolonged periods of infection and with extensive somatic hypermutation^1–3^. As the only protein expressed on the viral surface, Env, a heterotrimer of gp120 and gp41 subunits, has been the target of many vaccine efforts^4–8^. An efficacious vaccine regimen needs to prime and affinity mature bnAb responses against multiple conserved neutralizing epitopes on Env. The fusion peptide (FP) is critical for viral membrane fusion^9^ and consists of a conserved 15-20 amino acid long hydrophobic motif starting at the N-terminus of gp41 subunit of Env^10–15^. The FP neutralizing epitope is sterically occluded by 4 canonical glycans at positions N88, N230, N241 and N611^13^. FP is a viable target for HIV vaccine design efforts as potent bnAbs VRC34.01, PGT151, and ACS202 against the FP have been isolated from human elite neutralizers^11,16–18^.

Multiple vaccination efforts in non-human primates (NHP), rabbits and mice have elicited antibodies (Abs) that bind to the FP with a range of neutralizing and protective capabilities^10,13,15,19–23^. Consistent elicitation of FP-targeting Abs has been achieved by removal of specific canonical FP N-linked glycans in priming immunogens^13,23^. One such study immunized NHPs with BG505.SOSIP with three out of four canonical FP glycans removed except the glycan at residue N88^23^, a critical N-glycan for binding of FP-targeting bnAb VRC34.01^11^. The study showed that NHPs immunized with autologous trimers of increasingly native FP N-glycan presentation developed Abs that recognize the FP. Structural analysis of elicited polyclonal Ab responses, however, indicated that deletion of several glycans around the FP biased Ab recognition and maturation towards the glycan deletion at N611, thereby greatly limiting the potential for neutralization breadth and potency.

To overcome these limitations, we employed several immunofocusing strategies to our prime and boost immunogens to present the FP epitope in a more native-like context. We first immunized NHPs with recombinant BG505- and CH505-based chimeric immunogens either with full FP glycan occupancy (+N241) or with a single FP N-glycan deletion (ΔN241), to increase epitope accessibility in a more conservative manner than our prior approach^23^. Primed responses were first boosted with the BG505-CH505+N241 chimeric antigen and later boosted with a different trimer genotype with an identical FP to focus immune responses on the FP. Slow delivery immunization via osmotic pumps was employed for the priming phase to enhance generation of neutralizing antibodies (nAbs) as well as germinal center (GC) activity^20,24^. FP-specific GC B cell populations were observed over the course of the study as well as overall antigen specificity and memory B cell (MBC) population dynamics. Negative stain electron microscope polyclonal epitope mapping (nsEMPEM) was performed on animal plasma two or four weeks post immunization to map polyclonal Abs (pAbs) responses^25^. CryoEMPEM was later used to characterize pAbs at higher resolution to delineate residue level interactions with the immunogen to better comprehend the immunogenicity of the antigens used in the study^23,26^. Overall, while we did not observe robust FP priming and neutralization activity from this study, the techniques and tools used for polyclonal response evaluation have important implications for epitope-based vaccine research.

## Results

### FP Targeting Immunogen Design and Immunization regimen

Previous FP studies with non-glycosylated trimers around the FP have resulted in Ab responses biased towards the glycan deletions^13,23^. Instead, we hypothesized that a more conservative glycan deletion approach would induce FP-specific responses capable of broad HIV-1 coverage. In this study, we primed NHPs using chimeric trimers based on BG505 and CH505 with the V1, V2, V3, and V5 loops of CH505^27^, a clade C virus, grafted onto a clade A BG505 trimer background as wild-type BG505 does not carry two of the four canonical FP glycans at positions N241 and N230^28,29^. BG505 also lacks a conserved glycan at position 289, which opens up an epitope previously shown to be immunogenic, so we introduced a glycan at position 289 to mitigate the immunodominance of this glycan hole epitope (Fig 1a)^29^. In the experimental group, the chimeric priming immunogen presented three of the four FP glycans: N611, N88 and N230 but lacking in the N241 glycan (BG505-CH505ΔN241) (Fig 1a). Alternatively, the control group was primed with an autologous, chimeric immunogen with all four canonical FP glycans present, including N241 (BG505-CH505+N241). For the priming period, both groups of animals were given their respective soluble antigens and adjuvants via continuous delivery via an osmotic pump over a four-week period at a dose of 100 ug of antigen and 750 ug of SMNP adjuvant (Fig 1b)^23,24,30^. All subsequent immunizations were given as bolus with the same dosages. The animals were first boosted with the BG505-CH505+N241 immunogen at week 12 and later boosted with a heterologous, clade B trimer, AMC016^31^, but with BG505 FP sequence grafted, to further immunofocus on the FP and suppress boosting of pre-existing Abs against off-target epitopes (Fig 1b). The AMC016 trimer is highly glycosylated and therefore suitable to suppress BG505-CH505-specific off-target and glycan hole-directed pAb responses^29,32,33^. AMC016 has previously been shown to induce strain-specific, neutralizing Abs in rabbits after autologous immunizations^31^.

**Figure 1.**
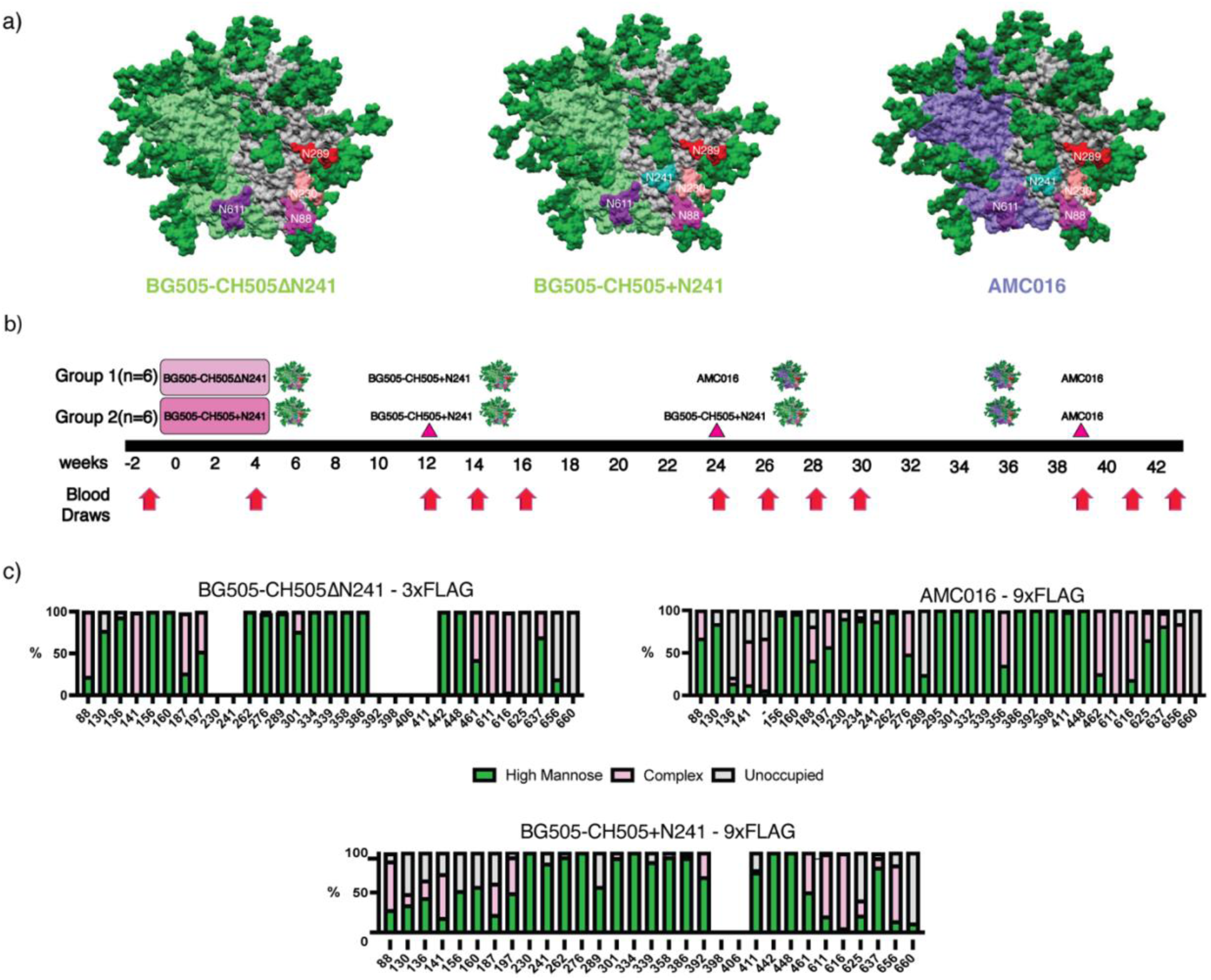
FP targeting Immunogens and Immunizations. a) Surface representations for BG505-CH505ΔN241 and BG505-CH505+N241, which are BG505 (Clade A)/CH505 (Clade C) chimeric trimers. The AMC016 trimer is a Clade B isolate. FP glycans are depicted as follows: N88 glycan in magenta, N230 glycan in salmon, N241 glycan in turquoise, and N611 glycan in purple. The N289 glycan is depicted in red. b) Immunization schedule. Osmotic pump immunogen delivery period is shaded in pink. Bolus immunizations are depicted with triangle indicator. c) Glycan Analysis of immunogens used in this study and impacts of Flag tag length on glycan occupancy.

To shield an otherwise immunodominant epitope at the base of soluble Env trimers, non-native N-glycans were introduced at positions 656 and 660 (Fig 1c) in all the constructs. Previous research has shown that membrane-bound trimers have higher glycan occupancy and improved glycan processing than soluble trimers due to longer retention in the ER/Golgi^34,35^. Here we employed an analogous strategy to extend the C-terminus of Env by introducing a Furin cleavage site (RRRRRR)^36^ after residue 664 of gp41, followed by a GS-rich linkage to a 3x-Flag-Tag (priming immunogens) or a 9x-Flag-Tag (boosting immunogens). This tag increases the translation time and is subsequently cleaved off by Furin protease and is therefore not present in the final immunogen (Fig S1a).

When a 3x-Flag-Tag was used, the introduced glycan at the C-terminus of gp41, N656 was less than 40% occupied while N660 remained completely unoccupied (Fig 1c). When a 9x-Flag-Tag was used, the glycan at N656 was ∼75% occupied, while glycan occupancy at N660 remained 0%. In these constructs there was also a large change in the glycan occupancy at canonical gp41 glycan at 625, which historically has low occupancy when expressed as a soluble trimer truncated at position 664 with a stop codon^37^. In the 3x-Flag-Tag constructs, the glycan at position 625 was completely unoccupied. However, in the autologous and heterologous boosting immunogens, which have 9x-Flag-Tags, there was a significant increase in glycan occupancy at position 625 to 40% in the BG505-CH505+N241 autologous boost and 98% in the AMC016 heterologous boost (Fig 1c). Although a PNGS sequon at N230 was present in the Env sequence for the BG505-CH505ΔN241 construct, mass spec analysis of this glycan site was unable to determine the occupancy levels of this FP glycan and so the occupancy of this site remains unknown.

Historically, BG505 SOSIP.664 has introduced cysteine residues at positions 501 of gp120 and 605 of gp41, forming an intra-protomer disulfide bond that stabilizes recombinant native-like Env trimers^28^. Despite this Env stabilization effort, antibody-mediated immunogen disassembly has been observed after repetitive autologous trimer immunizations^38^. Therefore, we introduced a new disulfide linkage in the BG505-CH505ΔN241 and BG505-CH505+N241 constructs, 501C-L663C, to prevent in-vivo trimer disassembly. Notably, this disulfide prevents binding of RM20A3, a base binding mAb isolated from macaques that were repeatedly immunized with BG505-based constructs (Figs S1b and S1c)^12,38,39^. The AMC016 construct has the cysteines at positions 501 of gp120 and 664 of gp41, which also prevent RM20A3 binding (Figs S1b and S1c).

### NsEMPEM detected FP response after heterologous boosting

Prior studies using EMPEM to map pAbs against different epitopes of the HIV Env protein provide a basis to evaluate the success of our immunofocusing approach ^12,26,40^. Here, we used nsEMPEM to analyze pAb responses at post immunization timepoints using probe antigens matched to the most recent immunogen (Fig 2a and Fig S2a)^25^. At week 4 only base binders were bound to the probing antigens, in this case the priming immunogens, in each group (Fig 2a and Fig S2a). This result aligns with previous observations showing base responses are consistently elicited early in soluble Env trimer immunizations in animal and human studies^12,23–26,40,41^.

**Figure 2.**
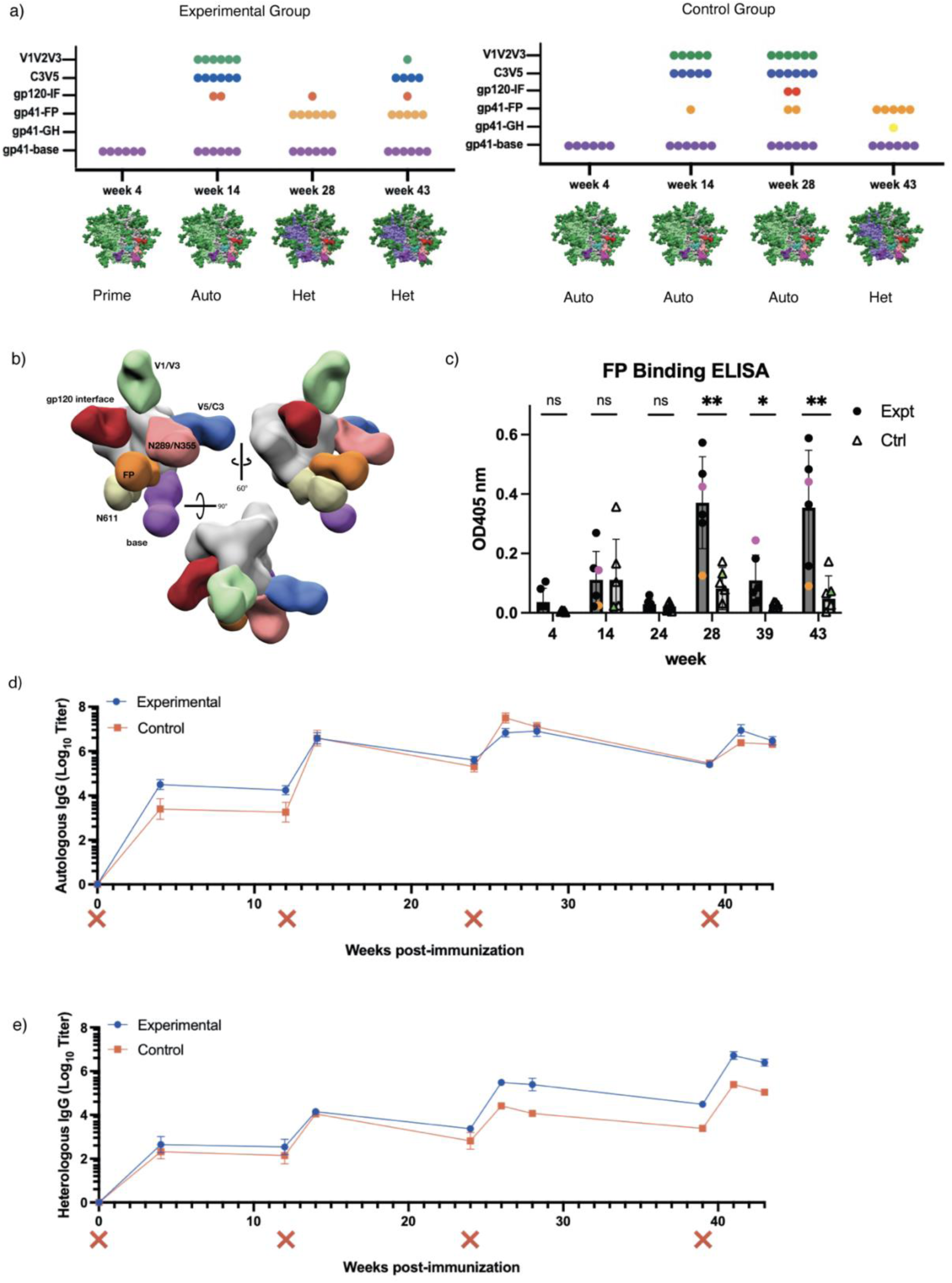
Humoral response analysis via ELISA and nsEMPEM. a) nsEMPEM dot plot results after each immunization b) EMPEM figure legend from multiple views c) FP-ELISA binding for time points also analyzed via nsEMPEM – experimental group (dark grey) versus control group (light grey). Animals RUu18, RQk18, and LJ66 are highlighted in green, pink and orange, respectively. d) Autologous trimer ELISA showing serum recognition of BG505-CH505+N241 trimer from control group (red) as well as experimental group (blue). (X) mark indicates immunization e) Heterologous trimer ELISA showing serum recognition of Heterologous AMC016 trimer from control group (red) as well as experimental group (blue)

After the initial priming immunizations, both groups were boosted with an autologous, chimeric immunogen with all four FP glycans present, BG505-CH505+N241, the same immunogen given to the control group for priming (Fig 1). Week 14 nsEMPEM analysis revealed more epitopes detected relative to week 4 (Fig 2a and Fig S2a). In addition to base binding pAbs, we observed pAbs bound to the C3/V5 region and V1/V2 region in most animals in both groups. One animal in the experimental group, LJ66, also developed a gp120/gp120 interface (IF) response (Fig S2a). One animal in the control group, RUu18, revealed an FP response (Fig S2a), however, follow-up cryoEM (discussed below) and FP-ELISA (Fig 2b, green triangle) demonstrated that the FP response observed in RUu18 via nsEMPEM did not specifically target the FP.

While FP pAbs were detected as early as six weeks in our earlier 2018 NHP study^23^, FP targeting pAbs were not observed via nsEMEPM until 4 weeks after the heterologous AMC016 trimer boosts at weeks 24 and 39 in the majority of animals within the experimental and control groups, respectively, in this study (Fig 2a and Fig S2a),. This suggests removal of only the N241 glycan around the epitope does not induce a robust priming immune response against the FP. Although FP responses were not observed at the earlier time points, FP-ELISA data suggested FP-recognizing Abs were elicited at low levels during chimeric BG505-CH505 construct immunizations (Fig 2c and Fig S2b).

Week 28 nsEMPEM analysis of the experimental group animals revealed base and FP responses, and one animal, LJ66, additionally showed a gp120/gp120 IF response (Fig 2a and Fig S2a). The control group, which had received three immunizations of the BG505-CH505+N241 antigen by this point, still exhibited base, V1/V2/V3, and V5 responses in all the animals. Two animals in the control groups revealed gp120/gp120 IF responses and FP responses, respectively, via nsEMPEM analysis. While our data suggests that a single canonical FP glycan deficiency on Env does not prime a robust FP response compared to a fully glycosylated trimer, the results from the heterologous AMC016 boost were encouraging.

To distinguish if post-AMC016 boost FP elicitation in the experimental group was due to epitope accessibility in the priming immunogens or introduction of the heterologous boosting immunogen, both groups were immunized a fourth and final time with the heterologous AMC016 trimer at week 39. NsEMEPM analysis of week 43 plasma samples in the control groups showed elicitation of FP-targeting Abs in five of the six control animals (Fig 2a). The experimental group, which had received two immunizations of the AMC016 trimer, elicited FP and base responses, as observed in week 28, however most of the animals now also exhibited AMC016-specific C3/V5 responses. Strain-specific responses after autologous immunizations but not after heterologous immunizations is a commonly observed phenomenon in multiple viral vaccination platforms^42,43^. Although nsEMPEM suggested both groups showed FP-directed responses after heterologous boosting, FP-ELISA showed that the experimental group had significantly higher recognition of the FP, suggesting that the N241 glycan hole in the BG505-CH505ΔN241 immunogen did impact FP-specific responses later boosted by the heterologous AMC016 trimer (Fig 2c and Fig S2b). Serum ELISAs showed negligible differences between experimental and control group antibody titers against the BG505-CH505+N241 probing antigen throughout the duration of the study (Fig 2d). However, the experimental group antibody titers had better AMC016 recognition after heterologous boosting at week 24, compared to the control group, even after the control group was heterologously boosted at week 39 (Fig 2e).

### Cross-reactive immunity observed via nsEMPEM

Cross-linking nsEMPEM of polyclonal sera enables detection of lower abundance antibodies^44^. Hence, prior to heterologous boosting we attempted this approach to reveal which epitopes might be boosted at weeks 24 and 39. The 2D class averages of samples cross-linked with the heterologous trimer suggests FP-recognizing Abs are present in plasma, which had not been observed in nsEMPEM analysis with BG505-CH505+N241 trimer (non-crosslinked) (Table S1). This suggestion was corroborated by the FP-ELISA data that showed FP-recognizing pAbs at the week 14 and 28 time points in both groups (Fig 2c). Thus, FP-directed Abs were elicited at low abundance and/or low affinity levels in the serum by the autologous immunogens. In all the animals, the 2D class averages also show the presence of a base-binding Ab that was able to cross-link with the heterologous immunogen prior to its immune exposure (Table S1). These observations suggest that cross-linked nsEMPEM analysis can reveal cross-linking epitopes between heterologous antigens and can be a valuable tool for vaccine design and boost immunogen selection for real time study evaluation^45^.

### Flow Cytometry Analysis Did Not Reveal Remarkable Changes in GC Populations

To gain visibility of the GC dynamics over the course of immunization, longitudinal lymph node fine needle aspirates (FNAs) were performed on the draining inguinal lymph nodes of each animal at several timepoints throughout the study (Fig S3). Cells collected during the FNAs were stained and analyzed using flow cytometry (Fig 3). Overall, GC B cell and TFH cell responses were observed in the sampled lymph nodes after initial priming and after the final boost (Fig 3a and 3b). GC response (as measured by GC B cell and TFH cell percentage) was largely unchanged after the boosts at weeks 12 and 24. Beginning at week 24, in addition to surface marker staining, FNA cells were also stained with a fluorescent probe of the heterologous boost immunogen. Interestingly, in the experimental group, the percentage of GC B cells collected from the lymph nodes that are positive for the heterologous AMC016 trimer are greater at week 24 than week 28, four weeks post-heterologous boost (Fig 3c). Despite no immunization with the AMC016 trimer, the control group exhibited a slight increase in heterologous AMC016 trimer recognition after BG505-CH505+N241 trimer boosts at week 24 (Fig 3c).

**Figure 3.**
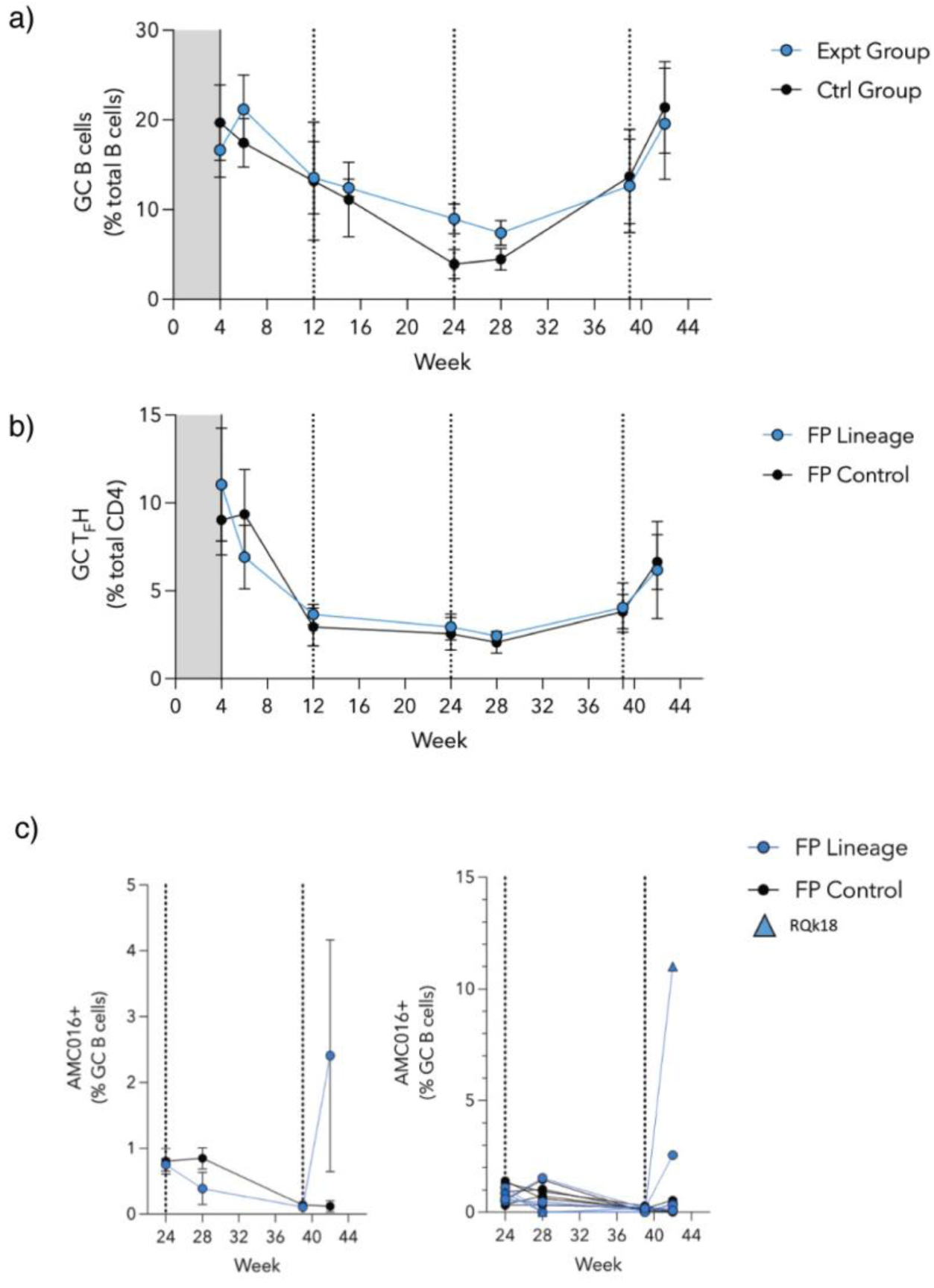
Flow cytometry reveals limited changes in GC B and TFH cell populations. a) Quantification of GC B cell kinetics as a percentage of total CD20+ B cells. Experimental group shown in blue circles while control group are shown in black circles. b) Quantification of GC Tfh cell kinetics as a percentage of total CD4+ T cells. c) % GC B cells that are AMC016+ following the boosts at weeks 24 and 39. Individual animals are shown on the graph to the right with animal RQk18 being the highest binder of the experimental group (depicted as a blue triangle).

### Serum Neutralization Detected After AMC016 Heterologous Boosting

We assessed heterologous or autologous neutralization of serum samples longitudinally using three pseudovirus (PV) neutralization panels: a PV panel sensitive to FP-targeting bnAbs, a BG505- and a CH505-specific mutant PV panel (Fig 4a, 4b, and 4c, respectively) (Tables S2, S3, and S4, respectively). The former is comprised of ten heterologous PVs selected for their neutralization sensitivity to FP-specific bnAbs^10,23^. Week 14 serum neutralization analysis revealed sporadic and weak neutralization for the FP sensitive panel, though slightly stronger against 3988.25 Tier 2 PV (Fig 4a; top panel). There was no detectable neutralization against BG505 and relevant mutant panel, except for two animals, one from each of the groups, that showed weak neutralization against the N611A mutant PV, which does not have a glycan present at position 611 of gp41 (Fig 4b; top panel). When tested against the CH505 PV panel, there was detectable neutralization against the Tier 2 transmitter/founder (TF) virus and against the Tier 1A w4.3 virus for the animals in both groups except 1 RM in the control group (Fig 4c).

**Figure 4.**
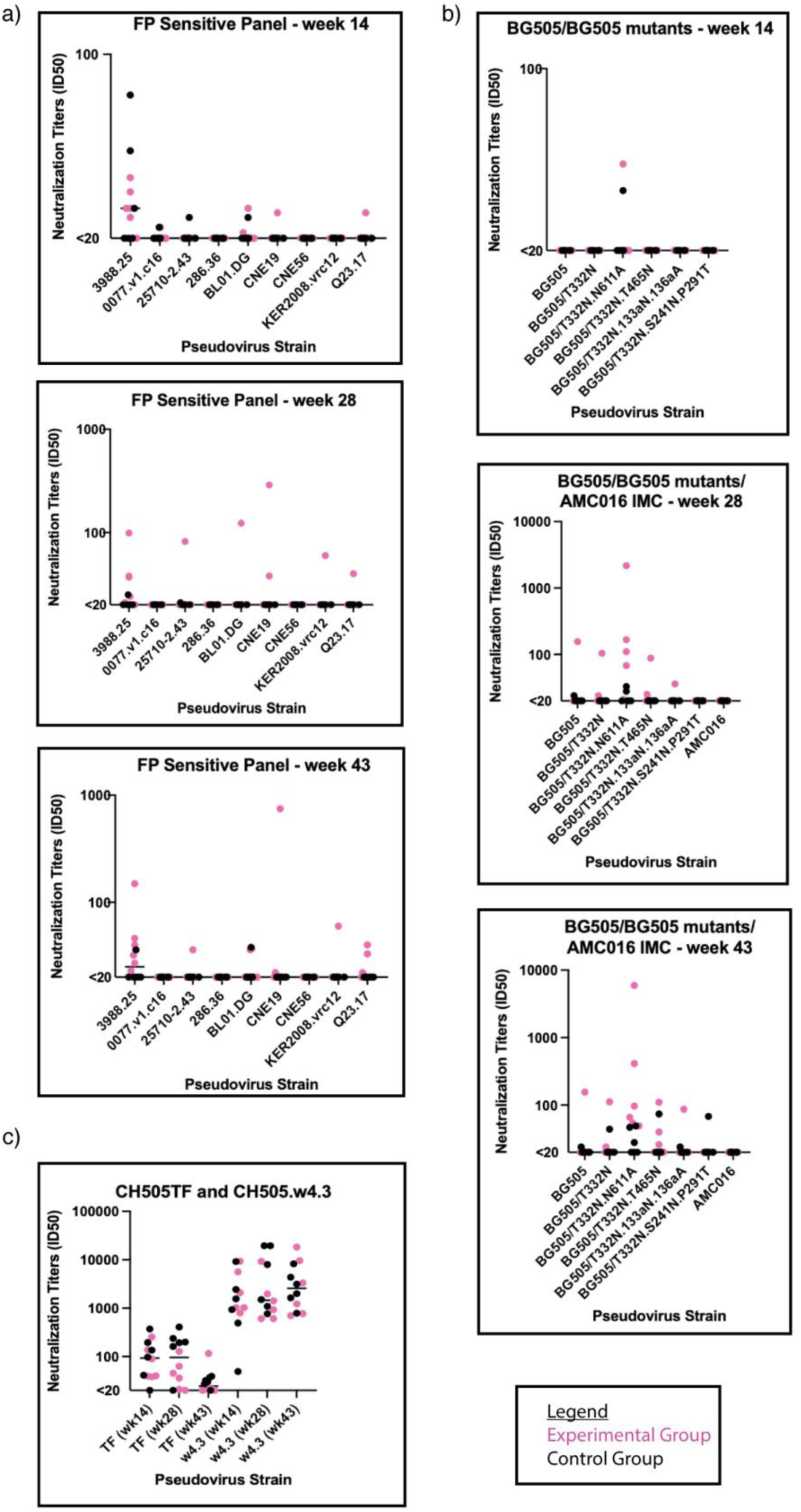
Serum neutralization sporadically observed after heterologous boosting. a) Comparison of FP-Sensitive pseudovirus panel serum neutralization between weeks 14 (top), 28 (middle), and 43 (bottom). (Experimental group is shown in magenta circles and the control group in black circles). Y-axis is in log scale. b) Comparison of BG505 pseudovirus panel serum neutralization between weeks 14 (top), 28 (middle), 43 (bottom). Y-axis is in log scale. c) Comparison of CH505 pseudovirus panel serum neutralization. Y-axis is in log scale.

Week 28 serum samples were analyzed for neutralization. One animal, RQk18, from the experimental group neutralized six of the nine PVs of the FP sensitive panel, all of which contained FP sequence identical to the FP sequence used in the immunogens of this study (Fig 4a; middle panel and Table S2). None of the control animals nor the rest of the experimental group developed cross-neutralizing capabilities for the viruses assessed in the FP panel except weak neutralization of Tier 2 3988.25 virus (Fig 4a; middle panel). Among the control group animals, none of them neutralized the BG505 PVs except weakly against the Tier 2 N611A PV seen in two of the animals (Fig 4b; middle panel). Within the experimental group, one animal, RQk18, neutralized five of six BG505 PV panel and four out of the six animals neutralized the N611A PV (Fig 4b; middle panel and Table S3). There was less serum neutralization of CH505 TF PV among the experimental group at this time point compared to the control group, which is expected since the experimental group received AMC016 trimers, and control received the BG505-CH505+N241 chimeric immunogen at week 24 (Fig 4c and Table S4).

After the week 39 boost with AMC016 trimer, FP responses were observed via nsEMPEM in both experimental and control groups in five out of six animals (Fig 2a). At week 43, one animal in the control group weakly neutralized one of nine PVs on the FP sensitive panel (Fig 4a; bottom panel and Table S2) and five of the six mutant BG505 panel (Fig 4b; bottom panel and Table S3). After this second AMC016 immunization, two more animals in the experimental group were able to neutralize additional PVs in the BG505 panel (Table S3) but animal RQk18 continued to weakly dominate in the FP sensitive panel neutralization with two other animals showing even weaker neutralization capacity (Table S2).

To test whether the weak serum neutralization profiles observed were mediated through FP recognition, we conducted an FP competition neutralization assay on a subset of PVs, whereby serum samples were titrated with FP-peptide, to capture any FP-recognizing antibodies in the serum (Fig S4 and Table S5). FP-peptide titration with VRC34.01 positive control showed significant decrease in PV neutralization, however, there was no impact on the serum neutralization profiles from immunized animals. The FP competition neutralization results suggest that the weak serum neutralization observed in a subset of the animals in the larger subset of PVs tested were due to non-FP directed neutralization.

### CryoEMPEM analysis reveals high resolution on-target and off-target immune responses

To better understand the types of FP region responses that were elicited in the animals and overall immunogenicity of our constructs, we conducted cryoEMPEM analysis^23,26^ on two animals - RQk18 at week 43 and RUu18 at week 14 time points (Fig S5 and Table S6) - to define the molecular interactions between antigen and pAb. Animal RQk18 developed weak neutralization breadth among the heterologous FP-sensitive PV panel after week 24 boosts, which persisted through the week 39 boosts. CryoEMPEM analysis of RQk18 week 43 fab response complexed with the heterologous AMC016 trimer resulted in three high-resolution maps: 2 define distinct pAb classes targeting the base of the trimer and one shows a pAb bound to the FP (Fig S5a). Animal RUu18 exhibited what appeared to be an “FP” response via nsEMPEM at week 14 despite FP-ELISA indicating poor FP recognition among the polyclonal IgG responses (Fig 2c, green triangle and Fig S2a). Multiple classes of pAbs were classified that targeted off-target BG505-CH505 epitopes: the base, V1V2V3 region and the non-specific “FP” (Fig S5b).

### RQk18 week 43 - FP

Animal RQk18 elicited a heterologous, neutralizing Ab response after week 24 AMC016 trimer boost, as determined by neutralization assays (Fig 4). To determine whether we could interpret high-resolution structural data and infer sequence information of the bound, polyclonal Abs, we turned to cryoEMPEM, which revealed two epitopes targeted after boosting with the heterologous trimer - the FP and base (Fig S5a). We resolved a 3.1 Å map with well-resolved density for the FP epitope as well as the variable region of the pAb, RQk-FP-A (Fig 5a).

**Figure 5.**
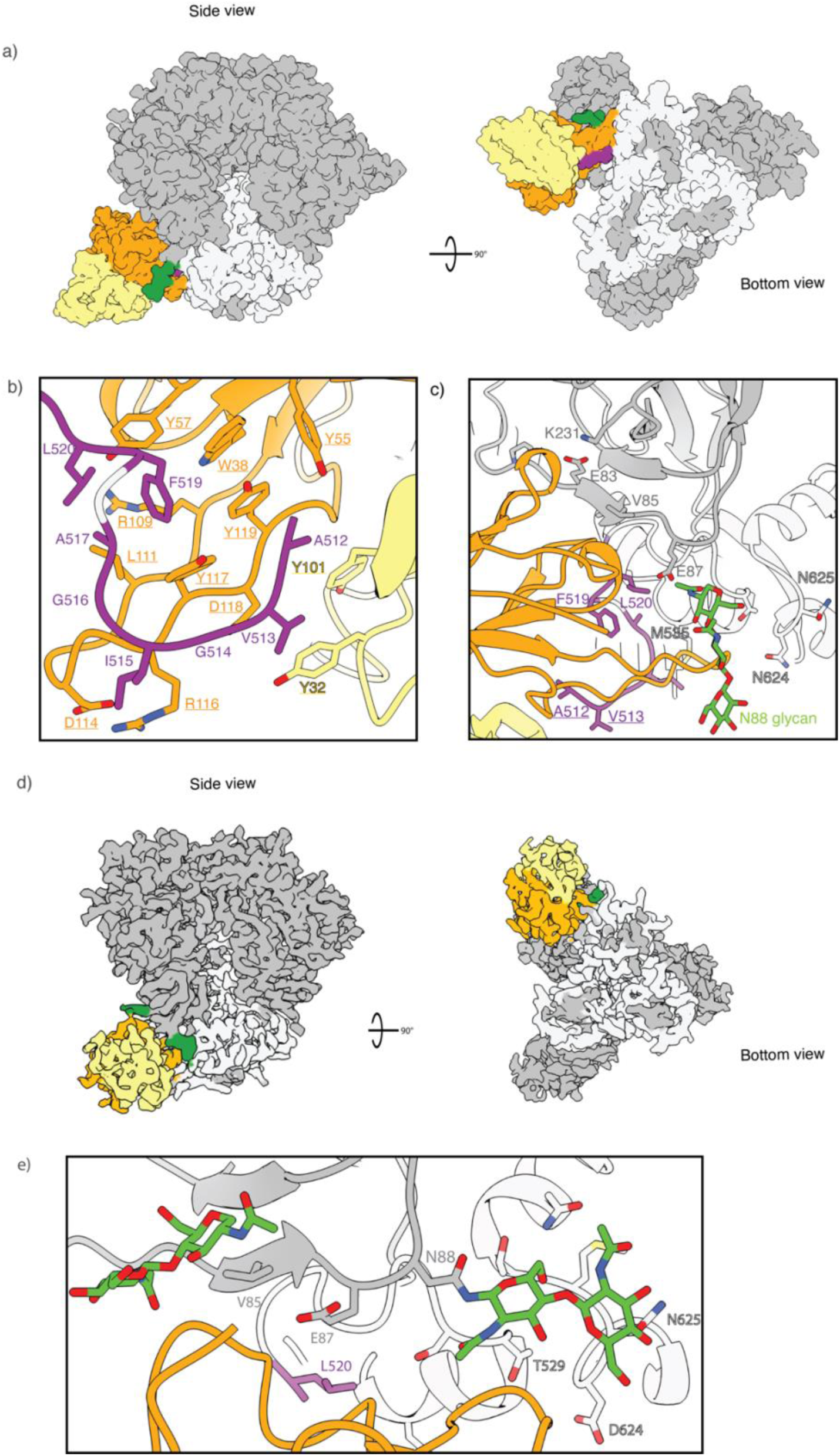
FP-targeting versus FP-proximal immune responses share overlapping epitopes. a) RQk-FP-A pAb1 from week 43 in complex with Heterologous Boost (left panel– side view; right panel – bottom view). The heavy chain (HC) is depicted in orange while the light chain (LC) is shown in yellow. The FP is shown in purple, the N88 glycan in green, gp120 subunit in grey and the gp41 subunit in white. b) RQk-FP-A pAb inferred hydrophobic and aromatic residues along its HCDR3 and LCDR1 and LCDR3 form a hydrophobic pocket that stabilizes the N-terminus of the FP c) RQk-FP-A pAb interacts with the N88 glycan and C1/C2 beta strands as well as the HR2 domain of gp41. d) RUu-FP-1 cryoEM map (Left – bottom view; right – side view) e) RUu-FP-1 pAb makes contacts with the C1 beta strand, HR2 and canonical FP glycans N241 and N88

The structural analysis revealed a fully resolved N-terminus of the FP, stabilized by aromatic side chains on the heavy and light chains of the Ab (Fig 5b). The HCDR3 of the heavy chain had a high abundance of aromatic residues, which help to stabilize the FP, a highly hydrophobic region (Fig 5b). This hydrophobic string of aromatic residues in the HCDR3 feature has been observed in two other bnAbs that target the FP - PGT151 and ACS202^18,46^. Such evidence indicates that NHPs are capable of eliciting FP Abs that have similar features to FP-directed human bnAbs. Apart from the HCDR3, the heavy chain contacts to the trimer include the CDR1 and CDR2 loops and a few residues of the framework region (FWR) 3 region. The light chain helps stabilize the N-terminus of the FP via CDRL3 loop interactions. Contacts with canonical FP Ab contacts as mentioned earlier were also observed for this pAb (Fig 5c).

### RUu18 week 14 - C1/C2 non-specific “FP” adjacent responses

Prior structural analysis of neutralizing, FP-targeting Abs have shown that the N-terminus of the FP is not typically resolved unless stabilized by an Ab^10,11,13,23^. NsEMPEM suggested animal RUu18 mounted an FP response by week 14 of the study, but an FP-specific ELISA suggested little FP interaction (Fig S2a and Fig 2c, green triangle). This indicated that there were immunogenic residues near the FP epitope that could not be resolved from true FP responses at low resolution. At high resolution, the “FP” Ab did not show engagement with the N-terminus of the FP (Fig 5d). Rather, the first FP residue observed interacting with RUu-FP-1 pAb is L520 (Fig 5e). The pAb instead predominantly interacted with the following residues: E83 main chain (99.5% global prevalence), I84 (45.21% prevalence), V85 (39.04% global prevalence), E87 (56.06% global prevalence), K231 (81.26% global prevalence), M535 (13.72% global prevalence). Previous epitope analysis of anti-FP bnAbs has shown that these non-FP residues commonly interact with anti-FP Abs^23^, consistent with the high overlap with the FP epitope despite lack of specific recognition of the FP N-terminus. Although this response was not visualized for the other control group members at week 14, FP-ELISA titers and nsEMPEM of week 43 results for the control group suggests cross-reactivity of C1/C2-partial FP response within all the control groups members and one of the experimental groups (LJ66) (Figs 2c and S2).

### RUu18 week 14 - V1V2V3

Three different V1V2V3-targeted pAbs maps were resolved to ∼4 Å from cryoEMPEM analysis of RUu18 at week 14 (Fig S6). Although this Ab response was not observed at the week 4 nsEMPEM analysis in either group, complexing polyclonal fabs isolated from week 14 with the BG505-CH505ΔN241 antigen showed binding of a V1V2V3 Ab to the priming trimer. Glycan analysis of the immunogens revealed an under occupancy of conserved glycans near the variable loops of gp120 of the boosting immunogen used at the week 12 boost for both groups (Fig 1c). Two of the three maps revealed a hydrophobic pocket that was created by hydrophobic residues in the substituted V1, V2, and V3 loops (Fig S6b). The HCDR3 of the Ab wedges between the loops to interact with a hydrophobic pocket formed by residues: A134, A136, I142, L175, V323 and I326. However, these maps also revealed the lack of glycosylation at a highly conserved, PNGS at residue N156 of the trimer-associated mannose patch (TAMP) (Fig S6c)^47^. Another 3.9 Å map, RUu-V1V2V3-5, showcases the HCDR3 of another Ab interaction with the variable loop hydrophobic region mentioned above, but in this case the presence of the N156 glycan would clash with the HCRD1 of this Ab instead of the light chain as in RUu-V1V2V3-2 (Fig S6d and S6c, respectively). Although the glycan sub-occupancy observed was unforeseen, the glycan hole and hydrophobic residues, synergistically, make this V1V2V3 region immunogenic and potentially distract from on-target responses.

### RUu18 week 14 - Base

A 4.1 Å map of a base Ab, RUu-Base-4 revealed an epitope that targeted the tryptophan clasp via HCDR1 and HDCR2 loops with long HCDR3 interactions with the N-terminus of gp120 (Fig S7a (left panel) and S7b). This epitope response was surprising as both autologous and heterologous immunogens in this study were shown to not bind RM20A3 (Figs S7a (middle panel) and S7c) via BLI analysis (Fig S1b). The lack of detectable RM20A3 binding initially led us to believe in the lack of exposure for this epitope in this construct, suggesting potential inaccessibility of this tryptophan clasp to immune recognition in-vivo. However, the adaptive immune system, unsurprisingly, adapted and targeted this neo-epitope and was able to accommodate the disulfide bond at the base of the trimer.

### RQk18 week 43 - Base

Two maps of AMC016 SOSIP bound by base Abs were also resolved during the analysis of RQk18 week 43 cryoEMPEM. A map with a base Ab, RQk-Base-C, was resolved to 3.1 A and also revealed an epitope that resembled the epitope of RM20A3 (Figs S7a, S7c and S7d)^12,48^. The epitope similarity between the two polyclonal complexes at this region suggests a conservation of the Ab response against this hydrophobic pocket at the base of the trimer (Fig S7a). A particularly well-resolved 2.9 Å map, RQk-Base-A, revealed an Ab that interacted heavily with the charged glutamic acid residues at the N-terminus of gp120 in AMC016 soluble trimer at positions E32, E32a, and E33 (Fig S7e). While we had hypothesized that the base of the trimer may be less targeted by antibodies with the introduction of the inter-protomer disulfide linkages, EMPEM results showed that the base of the trimer remained relatively immunodominant, although less susceptible to antibody induced disassembly due to the introduced disulfides.

### Structure To Sequence of RQk-FP

The quality of the 3.1 Å map of RQk-FP-A allowed for structure to sequence (STS) analysis to infer the amino acid sequence of the Ab based on the map characteristics and isolate FP-specific mAbs^49^. We were able to assign highest confidence scores for paratope residues and infer putative heavy- and light-chain sequences (Supp Data 01 and 02). PBMCs from week 42 for animal RQk18 were sequenced. The sequencing data and high resolution of the epitope-paratope interface enabled the design of four heavy chain and two light chain sequences that appeared to converge best with the highest confidence assignment regions. Sequencing analysis revealed the heavy chain to be of IGHV4-117*01 and IGHD2-12*01 allelic usage. Sequence analysis from week 14 PBMCs for animal RQk18 indicated that while IGHV4-117*01 and IGHD2-12*01 genes were used in different immunoglobulin (Ig) detected at this pre-heterologous boost time point, these genes were not paired in any of the 2170 clonotypes sequenced at this time point (Supp Data 03).

We designed a panel of eight monoclonal antibodies (mAbs) by pairing the four distinct heavy-chains with both light-chains and assessed binding kinetics against autologous and heterologous trimers, and subsequent nsEMPEM analysis (Tables S7-S9). Four out of eight mAbs showed detectable binding to the BG505-CH505+N241 and AMC016 Boost constructs, the majority of which showed higher affinity for BG505-CH505+N241 Boost (Fig 6a and S8a). Interestingly, FP-ELISA of the RQk-FP mAbs showed that the highest FP bindings mAbs showed reduced trimer recognition in the BLI results, suggesting these antibodies bound the FP productively the 3D spatial restraints within the trimer reduced FP binding (Fig 6b). Negative stain analysis of these trimer binding mAbs indicated that they did bind to the FP region of the trimer (Fig S8b). We undertook cryoEM with mAb 05, which had the best profile for affinity for the trimers (Fig 6a and S8a). A 3.9 Å map of RQk-FP-mAb-05 in complex with AMC016 and PGT122 (Fig 6c) showcases a similar binding mode to the pAb map (Fig 5A). When the model of pAb RQk-FP-A was docked in into the map of RQk-FP-mAb-05, there was a high degree of overlap of the FP antibody model and the FP mAb 05 density (Fig 6d). However, the N-terminus of the FP confirmation from pAb RQk-FP-A did not fit well into the FP density of RQk-FP-mAb-05 with the N-terminus not as clearly resolved (Fig 6e). The mAbs that showed binding to the immunogens via BLI were subjected to neutralization assays to determine whether the designed mAbs could recapitulate serum neutralization capacity. The mAbs were determined to be non-neutralizing (Fig 6f, Tables S10-S11), suggesting that the FP was not the neutralizing epitope seen in serum neutralization. However, the structure to sequence pipeline was still successful in discovering FP-specific mAbs without requiring expression and characterization of large panels of antibodies as has historically been done, showcasing the power and efficiency of this tool for epitope specific mAb isolation.

**Figure 6.**
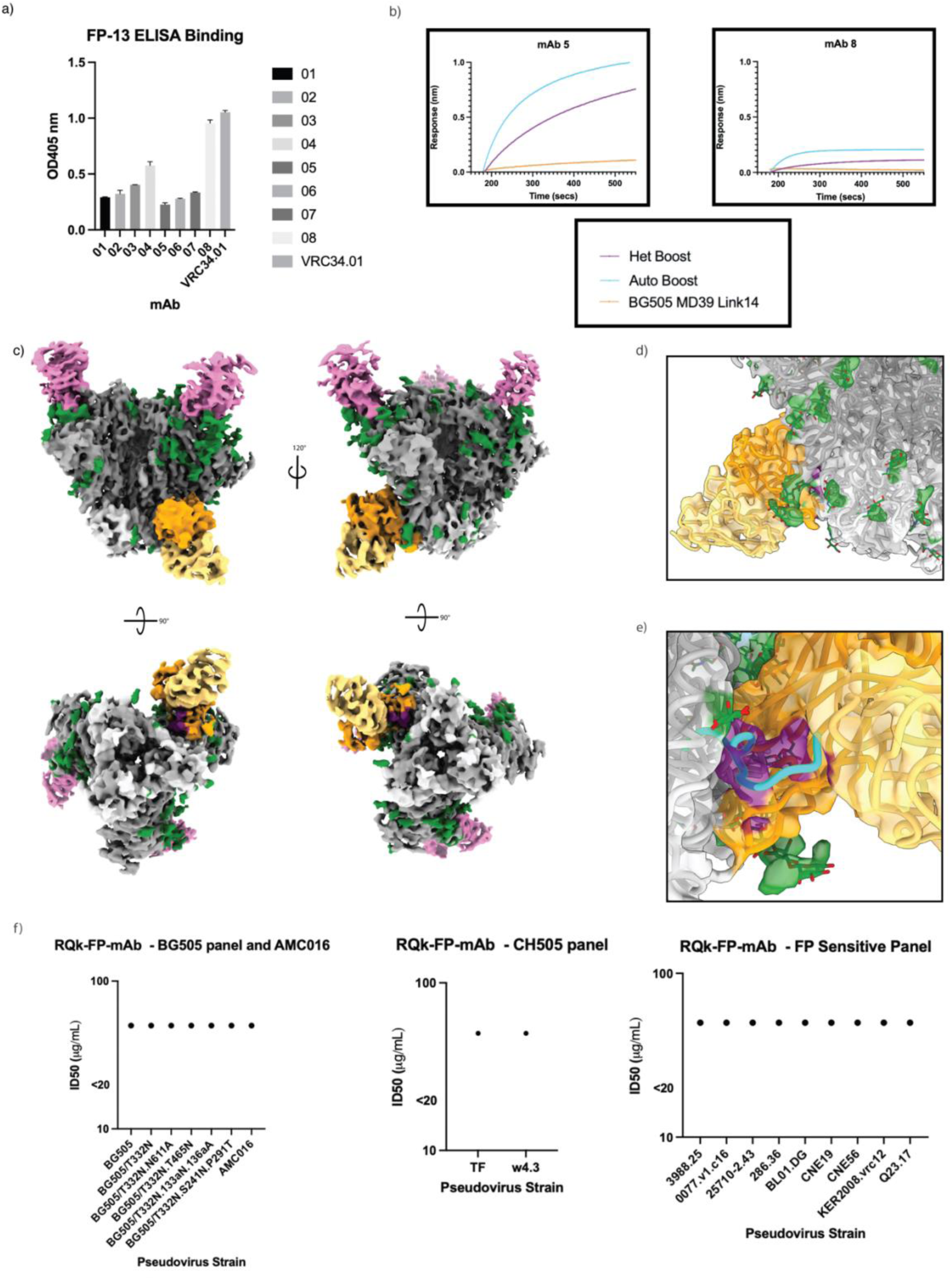
STS solves mAbs that bind to the FP region with varying FP interactions. a) FP-ELISA showcasing degree of FP interaction of the 8 different FP mAbs expressed from STS b) BLI curves of mAbs 05 (left) and 08 (right) interacting with trimers AMC016 (purple), BG505-CH505+N241 (blue) and negative control BG505 MD39 link 14 (orange) c) Colored EM maps of AMC016 (gp120 – dark gray, gp41 – light gray, glycans – green) in complex with RQk-FP-mAb 05 (heavy chain – orange, light chain – pale yellow), and PGT122 (pink). Bottom views showcase FP in purple. d) RQk-FPA pAb model, when docked into the mAb 05 EM map, shows high degree of overlap of FP antibody and FP density, shows same angle of approach and N88 glycan interactions e) RQk-FPA pAb modelled FP regions (blue) shows poor overlap with mAb 05 EM map density for FP region (purple), showing slightly different modes of FP engagement between the two antibodies f) Antibody neutralization panel of RQk-FP mAbs 01, 03, 05, and 07 show >50.00 μg/mL concentration for PV neutralization, which is interpreted as non-neutralizing. PV panels include BG505 and related mutants panel and AMC016 (right), a CH505 PV panel (middle) and a FP-sensitive PV panel (right)

## Discussion

In previous studies, NHPs immunized with HIV-1 Env soluble trimers with FP-proximal N-linked glycans deleted elicited FP-directed responses ^13,23^. Here, we report that having only one of the four canonical FP glycans (ΔN241) absent does not robustly prime FP-targeting Abs in NHPs. While the autologous BG505-CH505 immunizations primed low affinity and/or low abundance C1/C2 and FP B cell populations, only upon heterologous boosting were significant increases in C1/C2 and FP Abs observed via nsEMPEM and FP-ELISA titers, respectively. Heterologous boosting induced weak neutralization breadth in one animal and high resolution EMPEM analysis revealed a pAb that fully interacted with the FP. Our data supports previous findings that autologous boosters bolster strain-specific responses rather than eliciting desired cross-reactive immunity, which does occur with heterologous boosters^42,43,50,51^.

The increased FP-directed responses observed after delivery of heterologous boosters is unlikely to correlate with increased immunogenicity of the FP on the heterologous, AMC016 trimer. First, the FP motif in the BG505-CH505ΔN241, BG505-CH505+N241, and AMC016 immunogens are identical, an FP sequence grafted from BG505 (AVGIGAVF), as this is the most commonly circulating FP sequence amongst HIV isolates (www.hiv.lanl.gov). Repeated immunizations with BG505 SOSIP.664 trimer in rhesus macaques and rabbits also did not show consistent elicitation of FP directed Abs^8,11,19,32,52,53^, unless three or more glycans around the FP were removed in the trimer immunogens^13,23^. Secondly, rabbits immunized with wild-type AMC016 trimer, hence displaying the canonical FP sequence, AVGTIGAMF, did not elicit FP-directed Abs, suggesting weak immunogenicity of AMC016 SOSIP trimer when fully glycosylated^33^. Finally, the week 43 ELISA data suggest lack of FP recognition in the control group after heterologous boosting, which indicates FP on the heterologous trimer is not uniquely immunogenic in this study and did not prime a FP response.

Rather than priming a FP-directed response, the AMC016 immunogen boosted a small population of B cells targeting the conserved FP and C1/C2 regions that had been primed by the BG505-CH505ΔN241 or BG505-CH505+N241, respectively, in previous immunizations^54^. The FP-ELISA showed significant difference in FP recognition in the polyclonal response after heterologous boosting in the control group versus the experimental groups although both exhibit FP-like responses via nsEMPEM after AMC016 immunizations (Fig 2 and S2). High resolution cryoEMPEM reveals a newly characterized C1/C2 non-specific “FP” epitope in the control group that does not engage with the FP itself but C1/C2 residues that are commonly observed in FP specific epitopes as well^23^. The experimental group more consistently recognized the FP, suggesting that while the FP priming was not robust, the lack of N241 glycan in the experimental group priming immunogen did impact overall FP recognition in the immune response by the end of the study.

Lack of robust FP priming by the chimeric antigens limited the recall response breadth potential upon heterologous boosting ^55–59^. In the absence of high-affinity Ab competition in the serum, MBCs with moderate-to-low affinity for the boosting antigen can better form and populate recall germinal centers to expand and evolve these low affinity and/or low abundance, cross-reactive B cell clones with or without the occurrence of clonal bursts^42,45,54,60,61^. With little serum Ab competition, the low abundance FP- and C1/C2-targeting MBCs produced by the chimeric immunogens were able to engage the heterologous AMC016-based antigen. In some studies, the recall response would include lower affinity and/or lower abundance MBCs, differentiating into PCs to secrete Abs without further SHM, especially against heterologous viral variants^42,43,51,62^. Our results here indicate the preferential use of heterologous booster immunogens to focus the immune system on epitopes of interest; however, further research needs to be done to boost GC recall responses to drive iterative rounds of SHM to develop Ab breadth.

STS strategies were successful in pulling out epitope-specific monoclonal antibodies with varying levels of FP engagement. While they proved to be non-neutralizing, the antibodies constructed from our pAb EM map and sequencing data all recognized FP to different extents, which is quite exciting and impressive when compared with the historically laborious antibody discovery pipelines. More research has been done to improve the STS pipeline for improved isolation for target-specific antibodies, which will help advance the speed for bnAb discovery and target characterization as bnAbs are key for many vaccine efforts^63,64^.

High-resolution EM analysis reveals on-target (FP) and off-target (base, V1V2V3, C3/V5, and C1/C2 non-specific “FP”) epitope details of humoral immune recognition. Glycan sub-occupancy of the soluble trimer elicits off-target and distracting immune responses and more studies need to be conducted to optimize glycan occupancy and homogeneity in vaccine immunogens. The introduced disulfides at the base of the trimer, while unsuccessful in silencing base responses, were efficient in mitigating in vivo trimer disassembly that had been seen in prior soluble trimer immunizations^38^. The heterologous epitope boosting predictions would be a very valuable tool for vaccine development, to allow for immunofocusing towards epitopes of interest and away from immunogenic but nonprotective epitopes that consume valuable resources to elicit non-neutralizing and nonprotective immune responses. However, for a boosting immunogen to be successful, it will be imperative to design a priming immunogen strategy that strongly primes the desired Ab responses.

## Methods

### Immunogen and Probe Preparation

All immunogens and soluble trimeric, Avi-tagged probes were produced in HEK293F cells. Six days after transfection, the cell supernatants were harvested by centrifugation and purified using *Galanthus nivalis* lectin (Vector Laboratories) affinity chromatography followed by size exclusion chromatography (SEC) using a HiLoad 16/600 Superdex 200 pg (Cytiva) column. Further purification was performed using two rounds of negative selection columns produced using CNBr-activated Sepharose 4B beads (Cytiva) coupled to non-neutralizing mAbs according to the manufacturer’s instructions. First F105 negative selection was used to remove further monomeric, dimeric, and non-closed-state-conformation trimeric protein species and aggregates^65,66^. The unbound proteins, or closed state trimeric protein species or aggregates, in the F105 column flow through were then applied to a second 19b negative selection column to remove trimer species with a highly immunogenic conformation of the V3 glycan that leads to a lot of strain specific neutralizing Ab responses^67^. The flow through of the 19b negative selection column contained closed-state trimer protein species, which were then purified a second time using SEC. Immunogen preps were tested for endotoxin using an Endosafe instrument (Charles River). Avi-tagged proteins were then biotinylated following the manufacturer’s protocol for BirA enzyme (Avidity).

### Rhesus Macaques

Twelve rhesus macaques (Macaca mulatta) of Indian origin were acquired and housed at Emory National Primate Research Center and cared for according to NIH guidelines. Emory University Institutional Animal Care and Use Committee [IACUC# 201800298] approved this study. Animal care facilities are accredited by the U.S. Department of Agriculture (USDA) and the Association for Assessment and Accreditation of Laboratory Animal Care (AAALAC) International. Animals were treated with anesthesia (ketamine 5-10 mg/kg or telazol 3-6 mg/kg) and analgesics for procedures such as osmotic pump implantation and removal, subcutaneous immunization, blood draws, and lymph node fine needle aspirates as per veterinarian recommendations and IACUC approved protocols. When osmotic pumps were implanted, animals were kept in single, protected contact housing. At all other times, animals were kept in paired housing. Rhesus macaques were male, an age range of 3-4 years old, and at the start of the study a median weight of 5 kilograms. Animals were grouped to divide age, weight and gender as evenly as possible between the two groups. After completion of the proposed study, animals were transferred to other researchers upon the approval of the veterinarians.

### Animal Immunizations and sample collection

Osmotic pumps (Alzet model 2004) were loaded with 50 μg soluble Env trimer immunogen + 750 ug of SMNP Fig. 1B. Two pumps were subcutaneously (SC) implanted into each animal (one pump each in the left and right mid-thighs). The immunogen/adjuvant mixture was secreted continuously over the course of 4 weeks. The pumps were removed after 4 weeks. The animals were boosted at three timepoints s.c. with a bolus injection of 100 ug soluble trimer immunogens and 750 ug of SMNP (Figure 1) bilateral immunization split between the right and left mid-thigh. Blood was collected at various time points into CPT tubes for PBMC and plasma isolation. Serum was isolated using serum collection tubes and frozen. Plasma was used in ELISA and EMPEM analysis. Serum was used for neutralization assays.

### FP-ELISA

Streptavidin coated ELISA plates (Pierce Thermo Fisher), pre-blocked with BSA, were washed with wash buffer (1X TBS, 0.1% BSA, 0.5% Tween-20) 3x before coating with biotinylated FP-13 (InnoPep) (0.01 mg/mL in Wash Buffer) and left to rotate at 200 rpm and RT for two hours. Then plates were washed 3x before 100 μL of polyclonal IgG (1 mg/mL) for each sample timepoint or 100 μL RQk-FP mAbs 01-08 (150 μg/mL) was added to the plates and allowed to rotate at RT for an hour. Plates were then washed 3x before adding secondary antibody conjugated to alkaline phosphatase (AP) (anti-human Fab’2, goat antibody) and left to rotate for 30 minutes at RT. After washing 3x, developing solution is applied and allowed to develop while rotating for 30 minutes before stopping solution (2N NaOH) was used to stop the development. Plates were then read at 405 nm wavelength in a plate reader.

### Serum - ELISA

Serum samples from animals were serially diluted (starting at 1:80) and plated on pre-coated wells with BG505-CH505+N241 or AMC016 antigen or FP-13 (plates (Corning, Cat#: 3690) were coated with Streptavidin (Invitrogen, Cat#: 434302), blocked with 2% BSA and then incubated with biotinylated BG505-CH505+N241 or AMC016 or FP-13 probe). Env or FP binding-IgG antibodies were detected using HRP goat anti-human IgG (Jackson ImmunoResearch, Cat#: 109-035-098) and 1-Step™ Ultra TMB-ELISA Substrate Solution (Thermo Fisher Scientific, Cat#: 34029). The reaction was stopped with 2N of sulfuric acid (Ricca Chemical, Cat#: 8310-32) and read at 450 nm on a FlexStation 3 plate reader (Molecular Devices).

### TZM-bl cell-based neutralization assay

Env-pseudotyped virus neutralization assays completed at Duke were measured as a function of reductions in luciferase (Luc) reporter gene expression after a single round of infection in TZM-bl cells^68,69^. TZM-bl cells (also called JC57BL-13) were obtained from the NIH AIDS Research and Reference Reagent Program, as contributed by John Kappes and Xiaoyun Wu. Briefly, a pre-titrated dose of virus was incubated with serial dilutions of heat-inactivated (56 °C, 30 min) serum samples in duplicate for 1 h at 37 °C in 96-well flat-bottom culture plates, followed by addition of freshly trypsinized cells. One set of control wells received cells + virus (virus control) and another set received cells only (background control). After 48 h of incubation, cells were lysed and measured for luminescence using the Brit elite Luminescence Reporter Gene Assay System (PerkinElmer Life Sciences). ID50/IC50 and ID80/IC80 neutralization titers/concentrations are the dilution (serum/plasma samples) or concentration (mAbs) at which relative luminescence units (RLU) were reduced by 50% or 80% compared to virus control wells after subtraction of background RLUs from cells controls.

### FP Competition Neutralization Assay

TZM-bl neutralization assay using pseudoviruses CH505TF and BG505/T332N.N611A. Serum sample were titrated with or without FP-10, with a starting serum dilution of 1:20 and FP-10 concentration of 50 ng/mL; followed by 3-fold serial dilution for 8 total dilutions. VRC34.01 was titrated with or without FP-10, with a start concentration of 5 μg/mL and FP-10 concentration of 50 ng/mL; followed by 3-fold serial dilution for 8 total dilutions. Peptide and serum/mAb mixture were incubated at 37’C for 1 hr before viruses were added for routine neutralization assay. Cytotoxicity was observed in all wells with 50 ng/mL peptide (first dilution/concentration for serum/mAb titration). Serum dilution 1:10 and VRC34.01 concentration of 5 μg/mL were excluded from all assays for data analysis.

### LN-FNA and Cell Sorting

Lymph node fine needle aspirates were collected and processed as previously described^24^. B cells probes were prepared for flow cytometry by premixing individual probes with fluorochrome-conjugated streptavidin Ax647, BV421, or BV650 at RT for 20 minutes. For the FNA phenotyping panel, cells were incubated with appropriate conjugated probes in a stepwise fashion for 20 minutes at 4C. Following probe addition and incubation, surface antibodies (Table S12) were added directly to the cells and incubated for 30 minutes at 4C. Cells were washed twice, fixed for 30 minutes with 1% PFA at 4C, washed twice again, and acquired. Representative gating as well as antibody and clones for GC B cell and GC TFH cell flow cytometry is provided (Fig S2).

For the 10X B cell sorting panel, cells were incubated with appropriate conjugated probes for 30 minutes at 4C. Following probe addition and incubation, surface antibodies (Table S13) and appropriate TotalSeq cell hashing antibody were added directly to the cells and incubated for 30 minutes at 4C. Cells were then washed twice and acquired.

### Bulk B cell receptor sequencing

Total RNA was extracted from cryo-preserved peripheral blood mononuclear cells (PBMCs) of non-human primates (NHP) using the Qiagen RNeasy Kit (Catalog No. 75144), following the manufacturer’s protocol. The extracted RNA served as the template for reverse transcription, which was conducted using the Superscript IV Reverse Transcriptase and specific primers targeting the constant region of immunoglobulin heavy chains (IgM, IgG) and light chains (IgK, IgL) of *Macaca mulatta*. The cDNA was then subjected to a two-step PCR amplification process. In the first step, primers specific to the V-gene regions were used to amplify the target sequences. In the second step, the amplified products were further processed to include Illumina adapters necessary for high-throughput sequencing. The library preparation and PCR protocols were adapted from the methods described by Briney et al., 2019. Sequencing was performed on an Illumina NovaSeq 6000 platform using a paired-end 2×251 bp cycle run. The raw sequencing data was processed using abstar, which employed sequence merging, alignment, and annotation, producing a JSON-formatted output. The reference database used in the analysis contained unique amino acid sequences corresponding to the VDJ regions of heavy and light chain antibodies.

### nsEMPEM

Plasma samples were heat inactivated for 1 hour at 56C for live virus before purification via PrismA affinity column to isolate pAbs. pAbs were then digested to generate polyclonal fAbs with papain^70^. Once clean fabs were isolated, 500 ug of polyclonal fab were left to complexed with 15 ug of probing immunogen overnight at RT. Samples were then SEC purified to isolate trimer-fab complexes. After complex isolation, samples were stained and imaged on a FEI Talos microscope at a 73,000x magnification using Leginon. Relion v3.0 was used for 2D classification, 3D classification and 3D refinement. UCSF Chimera^71^ and Segger^72^ were used for visualizing and segmenting the EM density maps, respectively.

### CryoEMPEM sample preparation and imaging

∼10 mg of clean, polyclonal fab sample was complexed with 250 ug of probing trimer and left to complex overnight at RT. Samples were then SEC purified and the fab-trimer complexes were isolated. Samples were concentrated to 5-7 mg/ml. Just before sample application, samples were combined with lauryl maltose neopentyl glycol (LMNG) to a final concentration of 0.005. Samples were then vitrified using a Vitrobot mark IV (Thermo Fisher Scientific) on Quantifoil Cu 1.2/1.3 300C-mesh grids (Electron Microscopy Sciences) with 5-7 second blot times with a chamber set to 10C. Data collection for RUu18 week 14 was done on a 200 kV Talos Arctica and a K2 summit direct electron detector camera. Data collection for RQk18 week 43 was done on a 300 kV Titan Krios with a K2 summit direct electron detector camera.

### CryoEMPEM processing

CryoEMPEM was performed according to protocols previously described^26^. Micrograph movie frame alignments and dose-weighting was done through MotionCor2^73^. Data was processed using cryoSPARCv3.2.0^74^ and GCTF was used for CTF parameter estimation^75^. In cryosparc, template picker was used to pick trimer-fab complexes before running the particles through two rounds of 2D classification. Clean particles were selected for 3D homogenous refinement in C1 and C3 symmetries. Particles were then symmetry expanded for focused classification. In UCSF chimera, a 40 A mask was placed around 3 epitopes: FP, V1V2V3 and the Base. Symmetry expanded particles, with the C3 symmetry map and aligned sphere masks were used as inputs for 3D variability, done for each epitope investigated. Data processing for the different epitopes at this point were done separately but in parallel. Particles showing epitopes of interest were grouped and run through non-uniform refinement to obtain the high-resolution maps used for interpretation (Fig S3).

### CryoEMPEM model-building

An unliganded BG505 SOSIP (PDB 6V0R) was initially docked into each refined map in UCSF Chimera. Then, MODELLER was used to change the docked Env sequence to the autologous and heterologous boosts, for their respective data sets^76^. We docked polyalanine Fab models with fiducial markers (conserved disulfides and IMGT anchor residues) into each map and assigned the heavy and light chain orientations based on conformations of the FR2 and FR3 regions as well as CDR3 lengths. CDR lengths were determined and adjusted during manual model building using Coot^77,78^. Entire complexes were refined using both Rosetta^79^ and Phenix^80,81^.

### Structure to sequence

Structure to sequence predictions were done as previously described^49^. Inferred sequences were then searched within the B cell sequence library described above, which was incorporated into a Jupyter Notebook (www.jupyter.com) environment. Top hits were verified against the cryo-EM map to converge the pAb phenotype with the verified sequences. MolProbity^82^ and EMRinger^83^ were used to evaluate the final models before they were deposited to the PDB with polyalanine fab models representing the pAb Fv.

## Data Availability

CryoEMPEM densities and structures were deposited to the protein Data Bank (PDB) with the following accession codes: 9NHH, 9NHI, 9NHJ, 9NI9, 9NHK, 9NHL, 9NHM, 9NHN, 9NHO, and Electron Microscopy Data Bank (EMDB) with the following accession codes: EMD-49411 to EMD-49418, EMD-49457, and (FP-mAb05). Negative maps were deposited to EMDB with the following accession numbers: EMD-70702 - EMD-70717. Entries with polyclonal fab in complex with BG505-CH505ΔN241 are reported as BG505-CH505d241 due to character limitations for EMDB and PDB depositions.

## Acknowledgements

The authors thank Hannah L. Turner, Charles A. Bowman, Jean-Christophe Ducom, Bill Anderson, William Lessin and Anant Gharpure (The Scripps Research Institute) for electron microscopy, data acquisition and processing help. The authors also acknowledge Drs. Hailee Perrett and Lauren Holden for their help in manuscript preparation. This research was supported partly by the National Institute of Allergy and Infectious Disease of the National Institutes of Health Award Numbers AI145629 and AI136621. This work was funded by Cooperative Agreement award UM1 AI144462 in partnership with the Division of AIDS, NIAID. This research was funded in part by the Emory National Primate Research Center Grant Nos. ORIP/OD P51OD011132 and U42 PDP11023. The Emory National Primate Center is supported by the National Institutes of Health, Office of Research Infrastructure Programs/OD [P51OD011132 and U42 PDP11023] The authors are responsible for the contents of this manuscript, which do not necessarily represent the views of the US Government nor NIAID. The funders participated in no role of the study design, decisions, investigation, and preparation or publishing of this manuscript.

## Competing Interests

The authors declare no competing interests.

## Author Contributions

P.P.P. – Conceptualization, Data curation, Formal Analysis, Investigation, Methodology, Validation, Visualization, Writing – original draft, Writing – review & editing. C.A.C. – Conceptualization, Data curation, Formal Analysis, Investigation, Methodology, Validation, Writing – review and editing. J.Q. – Data curation, Formal Analysis, Investigation, Visualization, Writing – review & editing. D.G.C. – Resources, Writing – review & editing. D.B. – Formal Analysis, Writing – review & editing. A.S.T. – Investigation. C.A.E. – Resources. J.T.N. – Resources. S.T.R. – Investigation. H.G. – Investigation, Resources, Writing – review & editing. X.S. – Investigation, Resources, Writing – review & editing. K.M.G. – Investigation, Resources. J.H. – Investigation, Resources, Writing – review & editing. K.K.M. – Investigation, Resources, Writing – review & editing. E.B.A. – Resources. A.L. – Investigation, Resources, M.B.M. – Investigation, Resources, J.D.A. – Formal Analysis, Investigation, Visualization, Writing – review and editing. G.O. – Formal Analysis, Supervision. B.B. – Resources. M.C. – Resources. D.M. – Resources. G.S. – Resources. D.J.I. – Resources. S.C. – Conceptualization, Funding Acquisition, Project Administration, Resources, Supervision, Writing – review and editing. A.B.W. – Conceptualization, Funding Acquisition, Project Administration, Resources, Supervision, Writing – review & editing.

## Notes

### Competing Interest Statement

The authors have declared no competing interest.

### Summary of Updates

Additional experimental results testing Fusion Peptide binding of engineered antibodies designed from the reported structures, further assays testing polyclonal FP serum recognition and neutralization, and formal structure depositions codes from the PDB and EMDB.

