## Supplemental Material for "Immunofocusing on the conserved fusion peptide of HIV envelope glycoprotein in rhesus macaques"

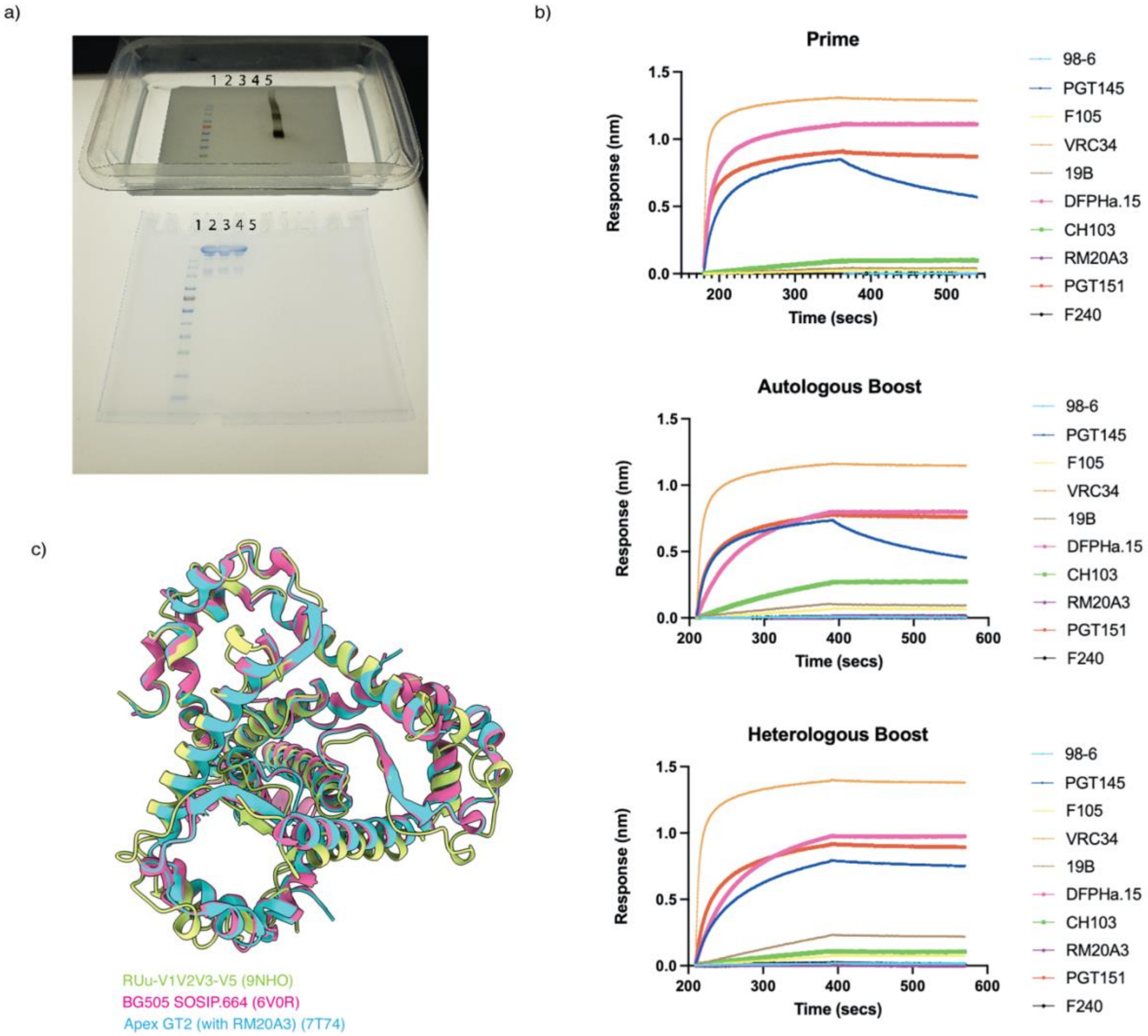

**Fig S1. Immunogen Design and Antibody Binding Profiles**

a) Anti-Flag Tag Western blot (top panel) and SDS-PAGE gel (bottom panel) of the immunogens with the same, following lanes: (1) Color Pre-stained Protein Standard, Broad Range (10-250 kDa) (NEB #P7719) (2) Prime Immunogen (3) Autologous Boost Immunogen (4) Heterologous Boost Immunogen (5) positive control recombinant Posi-tag epitope tag protein

b) BLI assessment of antigenicity against a panel of anti-Env mAbs: PGT145 (apex), VRC34 (FP), DFPHa.15 (FP), CH103 (CD4bs), RM20A3 (base), PGT151 (FP), F105 (anti-gp120, nnAb), 19b (V3), F240 (anti-gp41), 98-6 (anti-gp41).

c) Bottom view overlap of gp41 subunits of unliganded BG505 SOSIP.664 (501C-605C) in pink, Apex GT2 bound with RM20A3 (501C-605C) in blue and Autologous Boost cryoEMPEM map Uu-V1V2V3-5 (501C-663C) in green.

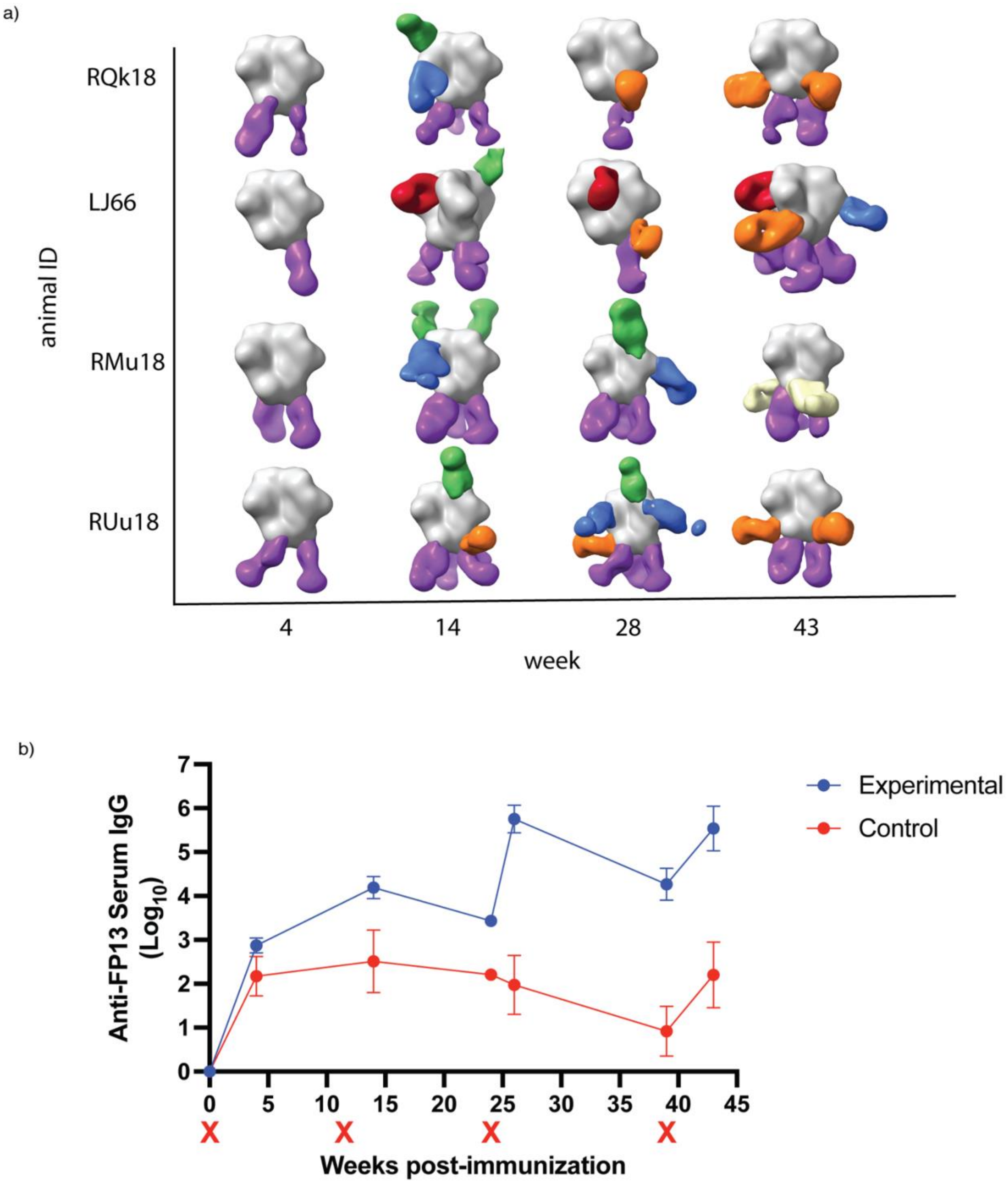

**Fig S2. nsEMPEM volumetric models and serum FP-ELISA**

a) Representative volumetric models of nsEMPEM analysis of two experimental group (RQk18 and LJ66) and two control group (RMu18 and RUu18) animals
b) Serum FP-ELISA using FP-13 as a probe to detect serum anti-FP levels with experimental group in blue and control group in red.

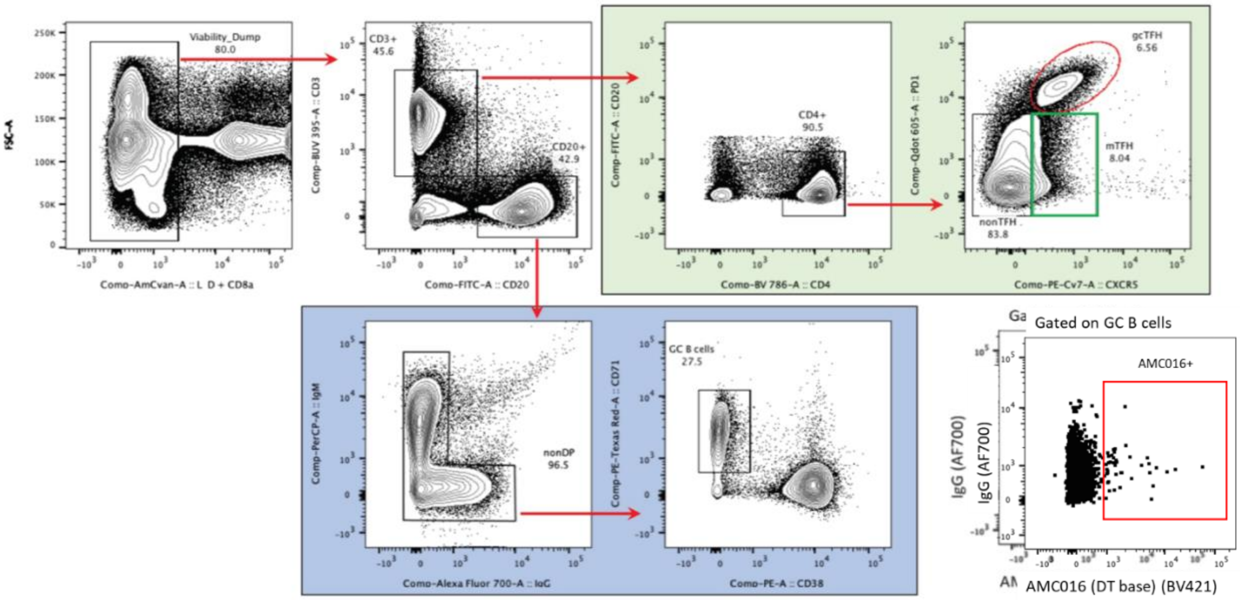

**Fig S3. Representative flow plots for FNAs**

Representative gating strategy and flow plots used for FNA analysis

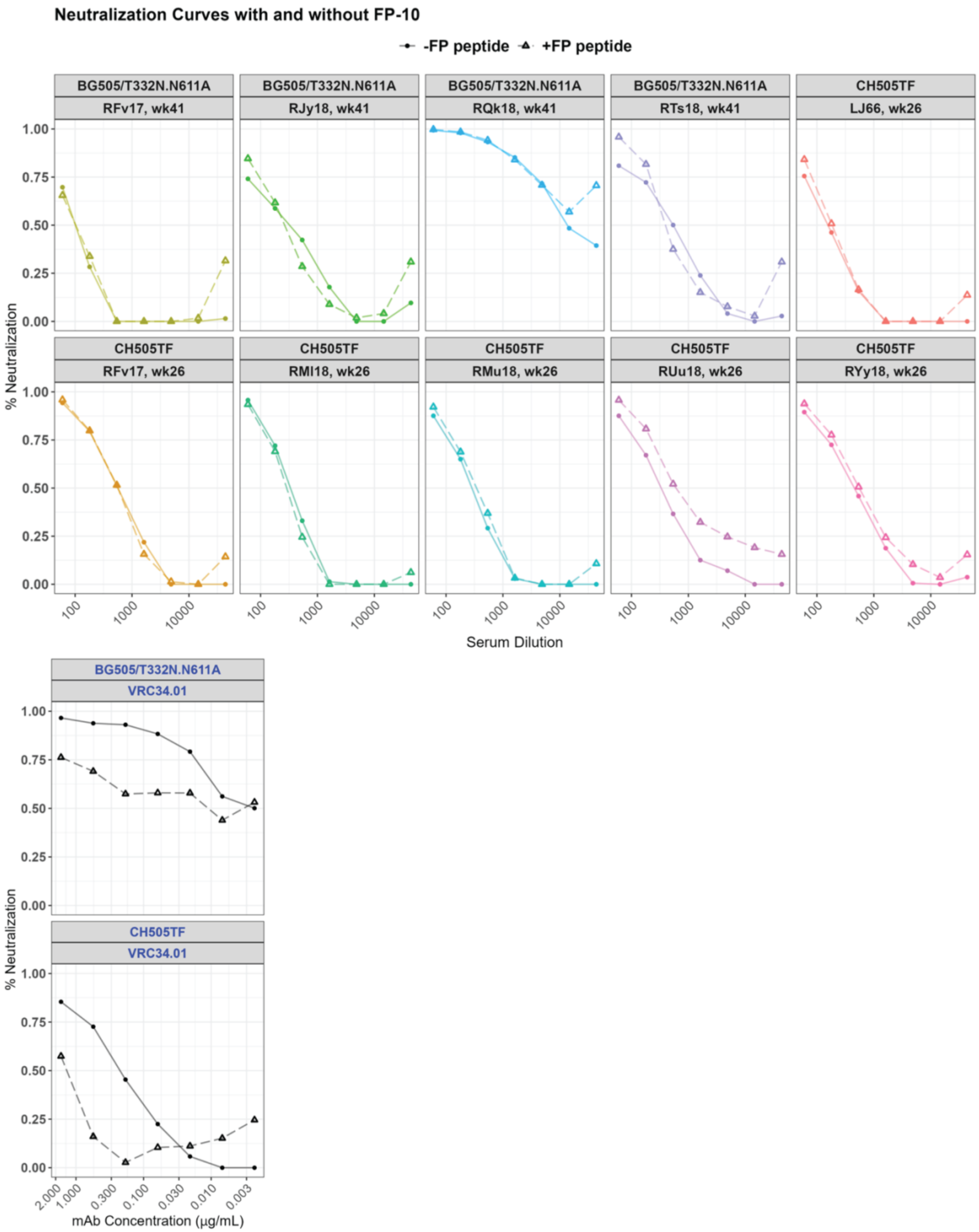

**Fig S4. FP Peptide Competition Neutralization Assays**

Serum neutralization assays with (dotted-lines) versus without (solid-lined) FP-10 spiked in to capture any FP-targeting antibody responses

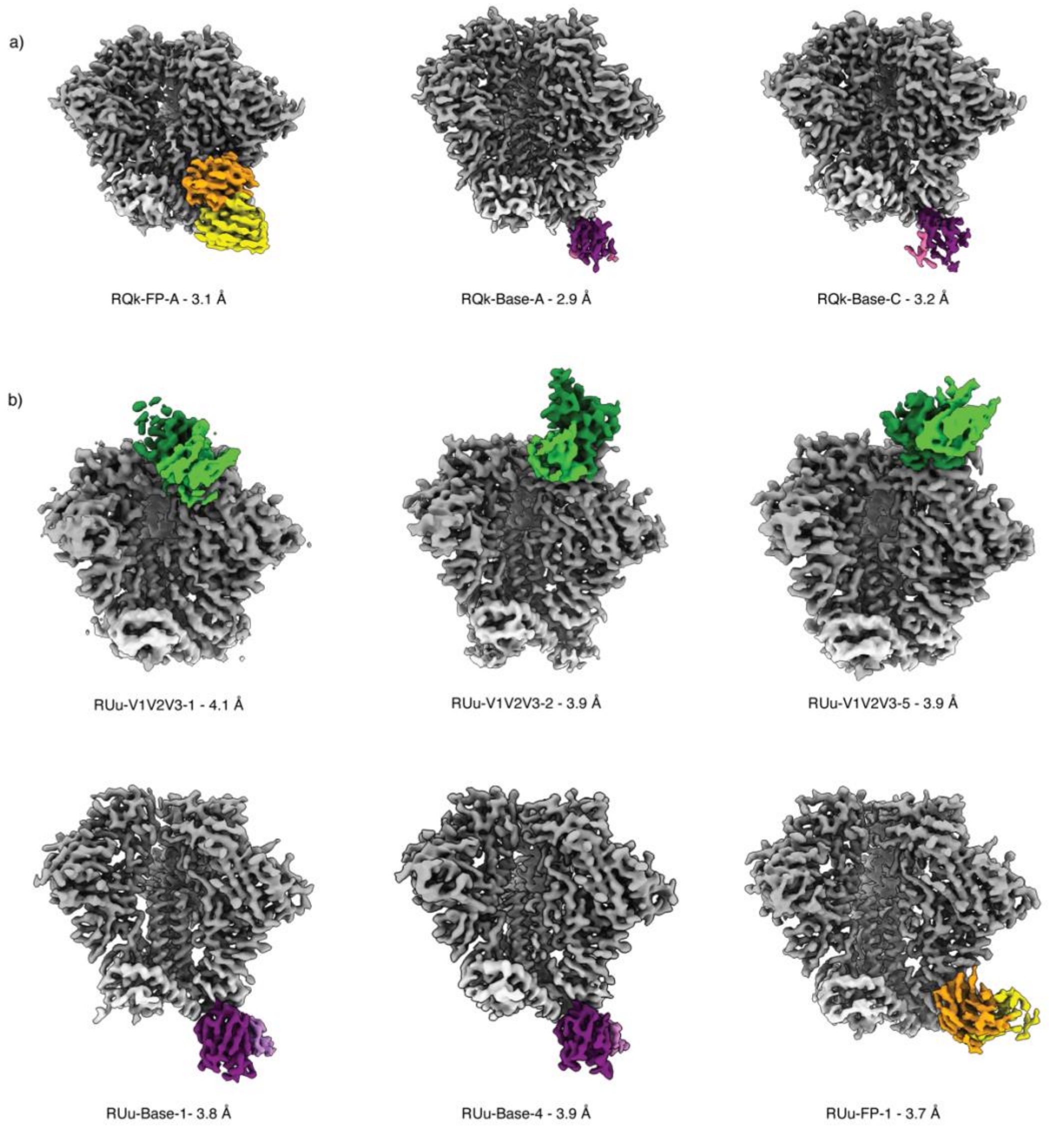

**Fig S5. CryoEMPEM High Resolution Maps**

a) Three high resolution maps were resolved from animal RQk18 week 43 polyclonal Fab response in complex with Heterologous Boost, including one against the FP and two maps against the base.
b) Six high resolution maps were resolved from animal RUu18 week 14 polyclonal Fab response in complex with Autologous Boost, including three against the V1V2V3 regions, two against the base and one against the C1/C2 non-FP-specific region.

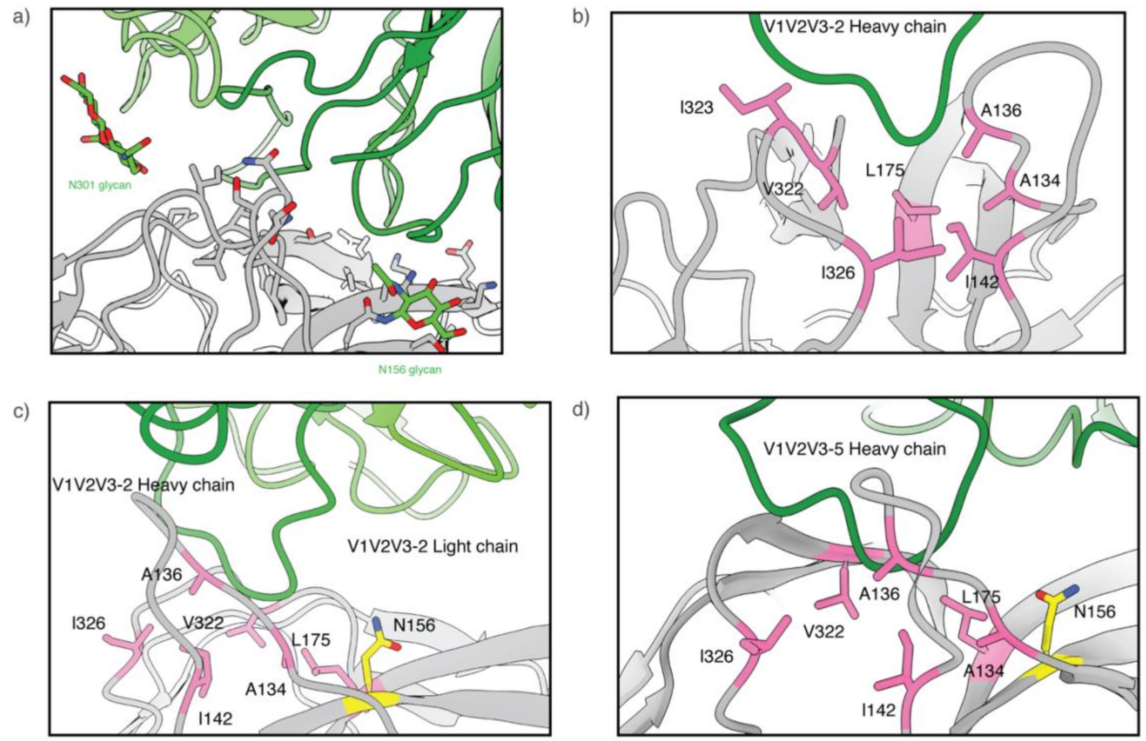

**Fig S6. V1V2V3 Off-Target Responses**

a) RUu-V1V2V3-1 (green) responses bracketed by N301 and N156 glycans
b) RUu-V1V2V3-2 view (green) 1 shows interactions with hydrophobic patch (pink)
c) RUu-V1V2V3-2 (green) view 2 shows lack of N156 glycan presence (yellow)
d) RUu-V1V2V3-5 (green) shows lack of N156 glycan presence (yellow) as well as hydrophobic patch (pink)

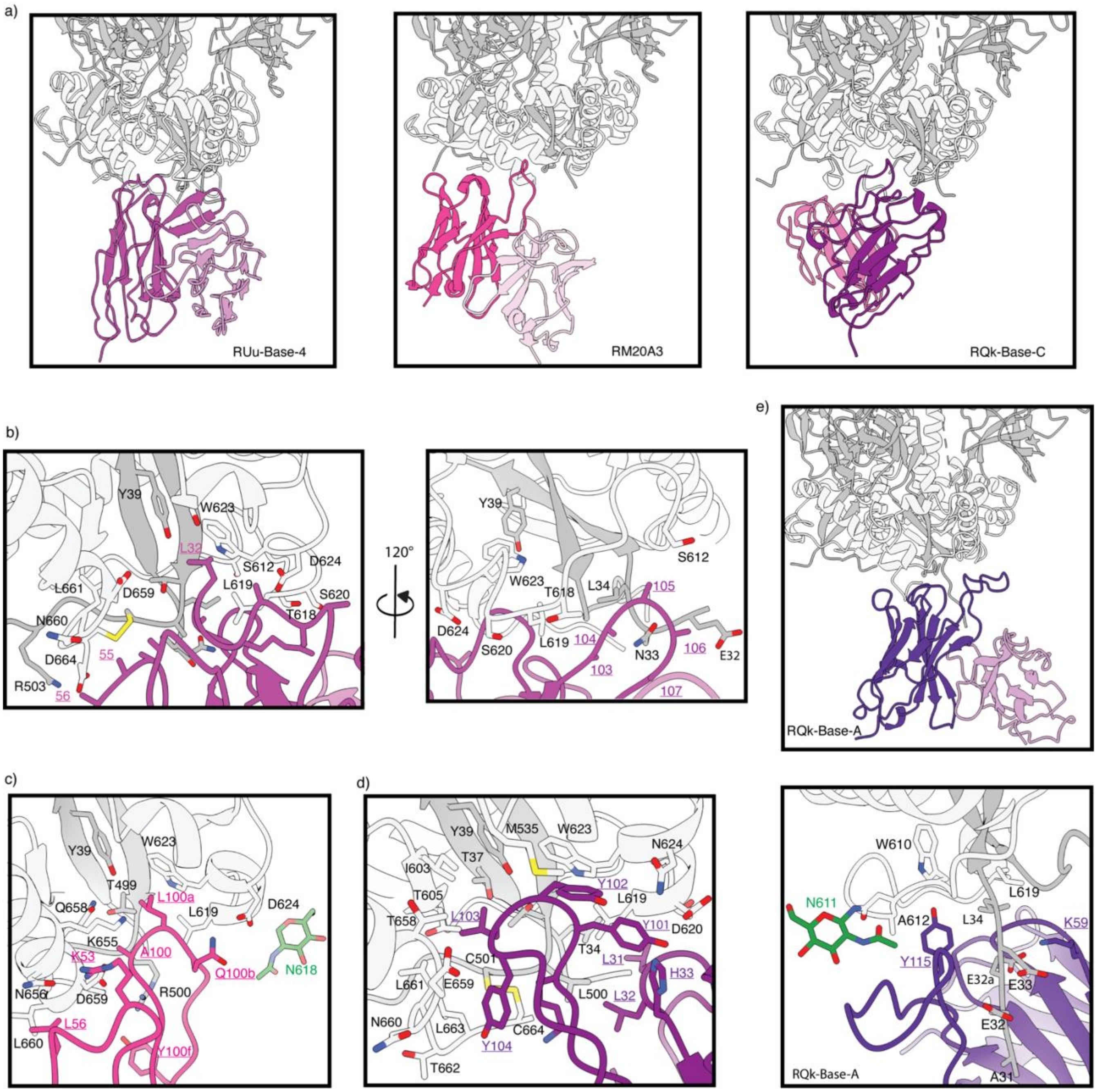

**Fig S7. Base Off-target Responses**

a) Base Antibodies target the tryptophan clasp (RUu-Base-4 – left panel; RM20A3 (PDB 7T74) – middle panel; RQk-BaseC – right panel)
b) RUu-Base-4 targets the tryptophan clasp with HCDR1 and HCDR2 (left panel) and N-terminus of gp120 with its HCDR3 (right panel)
c) RM20A3 (PDB 7T74) targets the tryptophan clasp with its HCDR3 interactions
d) RQk-Base-C targets tryptophan clasp with its HCDR3 interactions
e) RQk-Base-A targets the N-terminus of gp120 (top panel) with HCDR3 interactions with the charged residues at the N-terminus (bottom panel)

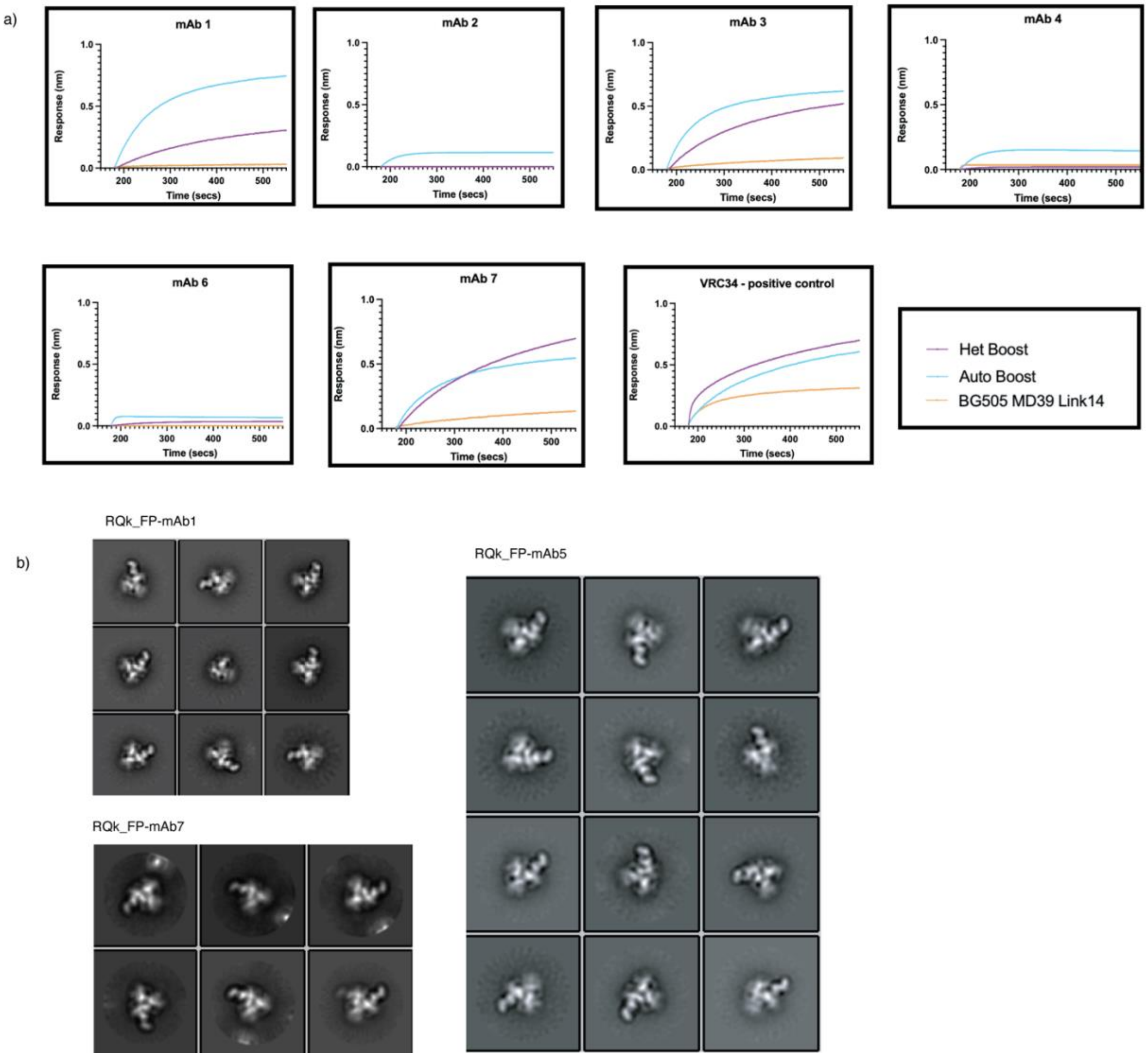

**Fig S8. SFS RQk_FP mAb BLI Binding Analysis and nsEM showcase FP and trimer interacting antibodies**

a) BLI Binding Analysis for recombinant expressed RQk_FP_mAbs and VRC34 (positive control) to the Heterologous Boost (purple), Autologous Boost (blue) and BG505 MD39 Link14 that does not present FP in a native confirmation due to a linked between C-terminus of gp120 and N-terminus of gp41.

b) ) nsEM 2D Classes of RQk_FP_mAbs for mAbs 01, 05, and 07

**Table S1. Cross-linked nsEMPEM**

| Animal ID (Group) | Time point (wk) | Probing Immunogen | 2D Classes |
|---|---|---|---|
| LJ66 (1) | 14 | AMC016 | 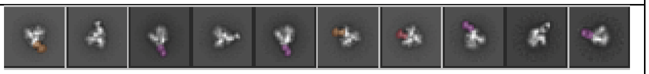 |
| RQk18 (1) | 14 | AMC016 | 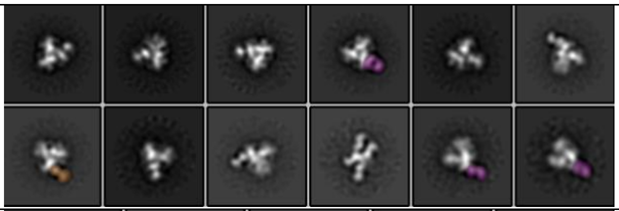 |
| RJy18 (1) | 14 | AMC016 | 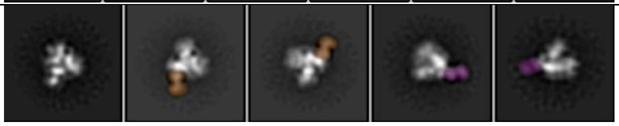 |
| RNp18 (1) | 14 | AMC016 | 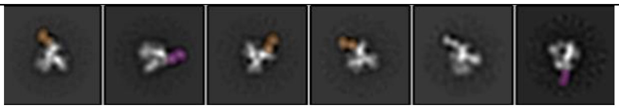 |

**Table S2. FP-sensitive pseudovirus neutralization data weeks 14, 28, 41 and 43.**

| Study Group# | Animal ID | Week | 25710-2.43 | 3988.25 | 0077.V1.C16 | CNE19 | CNE56 | KER2008.vrc12 | Q23.17 | 286.36 | BL01.DG |
|---|---|---|---|---|---|---|---|---|---|---|---|
| Control | RFv17 | 14 | <20 | <20 | <20 | <20 | <20 | <20 | <20 | <20 | <20 |
| Control | RTh18 | 14 | <20 | <20 | <20 | <20 | <20 | <20 | <20 | <20 | <20 |
| Control | RMl18 | 14 | <20 | **70** | <20 | <20 | <20 | <20 | <20 | <20 | <20 |
| Control | RMu18 | 14 | **24** | **26** | **22** | <20 | <20 | <20 | <20 | <20 | <20 |
| Control | RYy18 | 14 | <20 | **43** | <20 | <20 | <20 | <20 | <20 | <20 | <20 |
| Control | RUu18 | 14 | <20 | <20 | <20 | <20 | <20 | <20 | <20 | <20 | **24** |
| Experimental | LJ66 | 14 | <20 | **34** | <20 | <20 | <20 | <20 | **25** | <20 | **21** |
| Experimental | RNp18 | 14 | <20 | **30** | <20 | **25** | <20 | <20 | <20 | <20 | <20 |
| Experimental | RNz18 | 14 | <20 | <20 | <20 | <20 | <20 | <20 | <20 | <20 | <20 |
| Experimental | RJy18 | 14 | <20 | **26** | **22** | <20 | <20 | <20 | <20 | <20 | **26** |
| Experimental | RTs18 | 14 | <20 | **24** | <20 | <20 | <20 | <20 | <20 | <20 | <20 |
| Experimental | RQk18 | 14 | <20 | **26** | <20 | <20 | <20 | <20 | <20 | <20 | <20 |
| Control | RFv17 | 28 | <20 | <20 | <20 | <20 | <20 | <20 | <20 | <20 | <20 |
| Control | RTh18 | 28 | **21** | <20 | <20 | <20 | <20 | <20 | <20 | <20 | <20 |
| Control | RMl18 | 28 | <20 | <20 | <20 | <20 | <20 | <20 | <20 | <20 | <20 |
| Control | RMu18 | 28 | <20 | **25** | <20 | <20 | <20 | <20 | <20 | <20 | <20 |
| Control | RYy18 | 28 | <20 | <20 | <20 | <20 | <20 | <20 | <20 | <20 | <20 |
| Control | RUu18 | 28 | <20 | <20 | <20 | <20 | <20 | <20 | <20 | <20 | <20 |
| Experimental | LJ66 | 28 | <20 | **24** | <20 | **38** | <20 | <20 | <20 | <20 | <20 |
| Experimental | RNp18 | 28 | <20 | **37** | <20 | <20 | <20 | <20 | <20 | <20 | <20 |
| Experimental | RNz18 | 28 | <20 | <20 | <20 | <20 | <20 | <20 | <20 | <20 | <20 |
| Experimental | RJy18 | 28 | <20 | **21** | <20 | <20 | <20 | <20 | <20 | <20 | <20 |
| Experimental | RTs18 | 28 | <20 | **38** | <20 | <20 | <20 | <20 | <20 | <20 | <20 |
| Experimental | RQk18 | 28 | **82** | **99** | <20 | **289** | <20 | **60** | **40** | <20 | **123** |
| Control | RFv17 | 41 | <20 | <20 | <20 | <20 | <20 | <20 | <20 | <20 | <20 |
| Control | RTh18 | 41 | **78** | <20 | <20 | **31** | **162** | **34** | **54** | <20 | **63** |
| Control | RMl18 | 41 | <20 | <20 | <20 | <20 | <20 | <20 | <20 | <20 | <20 |
| Control | RMu18 | 41 | <20 | <20 | <20 | <20 | <20 | <20 | <20 | <20 | <20 |
| Control | RYy18 | 41 | <20 | <20 | <20 | <20 | <20 | <20 | <20 | <20 | <20 |
| Control | RUu18 | 41 | <20 | **33** | <20 | <20 | <20 | <20 | <20 | <20 | <20 |
| Experimental | LJ66 | 41 | **38** | **32** | <20 | **33** | **84** | **28** | **94** | <20 | **45** |
| Experimental | RNp18 | 41 | **23** | **32** | <20 | <20 | <20 | <20 | <20 | <20 | **26** |
| Experimental | RNz18 | 41 | <20 | **51** | <20 | <20 | <20 | <20 | <20 | <20 | <20 |
| Experimental | RJy18 | 41 | <20 | **21** | <20 | <20 | <20 | <20 | <20 | <20 | <20 |
| Experimental | RTs18 | 41 | **25** | **52** | <20 | **21** | <20 | <20 | **27** | <20 | **25** |

| | | | | | | | | | | | |
|---|---|---|---|---|---|---|---|---|---|---|---|
| Experimental | RQk18 | 41 | **46** | **177** | <20 | **754** | <20 | **29** | **37** | **22** | **34** |
| Control | RFv17 | 43 | <20 | <20 | <20 | <20 | <20 | <20 | <20 | <20 | <20 |
| Control | RTh18 | 43 | <20 | <20 | <20 | <20 | <20 | <20 | <20 | <20 | **38** |
| Control | RMl18 | 43 | <20 | <20 | <20 | <20 | <20 | <20 | <20 | <20 | <20 |
| Control | RMu18 | 43 | <20 | <20 | <20 | <20 | <20 | <20 | <20 | <20 | <20 |
| Control | RYy18 | 43 | <20 | <20 | <20 | <20 | <20 | <20 | <20 | <20 | <20 |
| Control | RUu18 | 43 | <20 | **36** | <20 | <20 | <20 | <20 | <20 | <20 | <20 |
| Experimental | LJ66 | 43 | <20 | **32** | <20 | <20 | <20 | <20 | **33** | <20 | **36** |
| Experimental | RNp18 | 43 | <20 | **27** | <20 | <20 | <20 | <20 | <20 | <20 | **20** |
| Experimental | RNz18 | 43 | <20 | **46** | <20 | <20 | <20 | <20 | <20 | <20 | <20 |
| Experimental | RJy18 | 43 | <20 | **23** | <20 | <20 | <20 | <20 | <20 | <20 | <20 |
| Experimental | RTs18 | 43 | <20 | **40** | <20 | **22** | <20 | <20 | **22** | <20 | <20 |
| Experimental | RQk18 | 43 | **36** | **149** | <20 | **747** | <20 | <20 | <20 | <20 | **37** |

Values are the serum dilution when relative luminescence units (RLUs) were reduced 50% when compared against virus control wells.

**Table S3. BG505 pseudovirus neutralization data weeks 14, 28, 41 and 43.**

| Study Group | Animal ID | Week | SVA-MLV | BG505 | BG505/T332N | BG505/T332N.N611A | BG505/T332N.T465N | BG505/T332N.133aN+136aA | BG505/T332N.S241N.P291T |
|---|---|---|---|---|---|---|---|---|---|
| Control | RFv17 | 14 | <20 | <20 | <20 | <20 | <20 | <20 | <20 |
| Control | RTh18 | 14 | <20 | <20 | <20 | <20 | <20 | <20 | <20 |
| Control | RMl18 | 14 | <20 | <20 | <20 | <20 | <20 | <20 | <20 |
| Control | RMu18 | 14 | <20 | <20 | <20 | <20 | <20 | <20 | <20 |
| Control | RYy18 | 14 | <20 | <20 | <20 | **34** | <20 | <20 | <20 |
| Control | RUu18 | 14 | <20 | <20 | <20 | <20 | <20 | <20 | <20 |
| Experimental | LJ66 | 14 | <20 | <20 | <20 | <20 | <20 | <20 | <20 |
| Experimental | RNp18 | 14 | <20 | <20 | <20 | <20 | <20 | <20 | <20 |
| Experimental | RNz18 | 14 | <20 | <20 | <20 | <20 | <20 | <20 | <20 |
| Experimental | RJy18 | 14 | <20 | <20 | <20 | **43** | <20 | <20 | <20 |
| Experimental | RTs18 | 14 | <20 | <20 | <20 | <20 | <20 | <20 | <20 |
| Experimental | RQk18 | 14 | <20 | <20 | <20 | <20 | <20 | <20 | <20 |
| Control | RFv17 | 28 | <20 | <20 | <20 | | <20 | <20 | <20 |
| Control | RTh18 | 28 | <20 | <20 | <20 | | <20 | <20 | <20 |
| Control | RMl18 | 28 | <20 | <20 | <20 | | <20 | <20 | <20 |
| Control | RMu18 | 28 | <20 | <20 | <20 | | <20 | <20 | <20 |
| Control | RYy18 | 28 | <20 | <20 | <20 | | <20 | <20 | <20 |
| Control | RUu18 | 28 | <20 | <20 | <20 | | <20 | <20 | <20 |
| Experimental | LJ66 | 28 | <20 | **24** | **24** | **68** | **25** | <20 | <20 |
| Experimental | RNp18 | 28 | <20 | <20 | <20 | **167** | <20 | <20 | <20 |
| Experimental | RNz18 | 28 | <20 | <20 | <20 | <20 | <20 | <20 | <20 |
| Experimental | RJy18 | 28 | <20 | <20 | <20 | **21** | <20 | <20 | <20 |
| Experimental | RTs18 | 28 | <20 | <20 | <20 | **110** | <20 | <20 | <20 |
| Experimental | RQk18 | 28 | <20 | **56** | **104** | **2160** | **88** | **36** | <20 |
| Control | RFv17 | 41 | <20 | <20 | <20 | **134** | <20 | <20 | <20 |
| Control | RTh18 | 41 | <20 | **37** | **83** | **42** | **90** | <20 | **61** |
| Control | RMl18 | 41 | <20 | <20 | <20 | <20 | <20 | <20 | <20 |
| Control | RMu18 | 41 | <20 | <20 | <20 | <20 | <20 | <20 | <20 |
| Control | RYy18 | 41 | <20 | <20 | <20 | **56** | <20 | <20 | <20 |
| Control | RUu18 | 41 | <20 | <20 | <20 | **25** | <20 | <20 | <20 |
| Experimental | LJ66 | 41 | <20 | **31** | **57** | **70** | **53** | **30** | **33** |
| Experimental | RNp18 | 41 | <20 | <20 | <20 | **77** | <20 | <20 | <20 |
| Experimental | RNz18 | 41 | <20 | <20 | <20 | **39** | <20 | <20 | <20 |
| Experimental | RJy18 | 41 | <20 | <20 | <20 | **226** | <20 | <20 | <20 |

| | | | | | | | | | |
|---|---|---|---|---|---|---|---|---|---|
| Experimental | RTs18 | 41 | <20 | **27** | **27** | **1,320** | **35** | **34** | <20 |
| Experimental | RQk18 | 41 | <20 | **221** | **189** | **8,766** | **174** | **178** | **34** |
| Control | RFv17 | 43 | <20 | <20 | <20 | **49** | <20 | <20 | <20 |
| Control | RTh18 | 43 | <20 | **24** | **44** | **28** | **74** | **24** | **68** |
| Control | RMl18 | 43 | <20 | <20 | <20 | <20 | <20 | <20 | <20 |
| Control | RMu18 | 43 | <20 | <20 | <20 | <20 | <20 | <20 | <20 |
| Control | RYy18 | 43 | <20 | <20 | <20 | **47** | <20 | <20 | <20 |
| Control | RUu18 | 43 | <20 | <20 | <20 | <20 | <20 | <20 | <20 |
| Experimental | LJ66 | 43 | <20 | **21** | **39** | **49** | **40** | <20 | <20 |
| Experimental | RNp18 | 43 | <20 | <20 | <20 | **54** | <20 | <20 | <20 |
| Experimental | RNz18 | 43 | <20 | <20 | <20 | **66** | <20 | <20 | <20 |
| Experimental | RJy18 | 43 | <20 | <20 | <20 | **97** | <20 | <20 | <20 |
| Experimental | RTs18 | 43 | <20 | <20 | <20 | **413** | **26** | <20 | <20 |
| Experimental | RQk18 | 43 | <20 | **156** | **112** | **5,928** | **110** | **87** | <20 |

Values are the serum dilution when relative luminescence units (RLUs) were reduced 50% when compared against virus control wells.

**Table S4. CH505 and AMC016 pseudovirus neutralization data weeks 14, 28, 41 and 43.**

| Study Group# | Animal ID | Week | CH505TF | CH505.w4.3 | AMC016 |
|---|---|---|---|---|---|
| Control | RFv17 | 14 | **97** | **2,434** | NA |
| Control | RTh18 | 14 | <20 | **49** | NA |
| Control | RMl18 | 14 | **41** | **492** | NA |
| Control | RMu18 | 14 | **195** | **931** | NA |
| Control | RYy18 | 14 | **370** | **9,200** | NA |
| Control | RUu18 | 14 | **135** | **1,551** | NA |
| Experimental | LJ66 | 14 | **253** | **5,636** | NA |
| Experimental | RNp18 | 14 | **39** | **1,027** | NA |
| Experimental | RNz18 | 14 | **137** | **9,286** | NA |
| Experimental | RJy18 | 14 | **89** | **2,086** | NA |
| Experimental | RTs18 | 14 | **40** | **1,027** | NA |
| Experimental | RQk18 | 14 | **38** | **795** | NA |
| Control | RFv17 | 28 | **408** | **19,513** | <20 |
| Control | RTh18 | 28 | <20 | **765** | <20 |
| Control | RMl18 | 28 | **199** | **1,095** | <20 |
| Control | RMu18 | 28 | **193** | **1,505** | <20 |
| Control | RYy18 | 28 | **162** | **19,463** | <20 |
| Control | RUu18 | 28 | **238** | **7,975** | <20 |
| Experimental | LJ66 | 28 | **128** | **9,264** | <20 |
| Experimental | RNp18 | 28 | <20 | **604** | <20 |
| Experimental | RNz18 | 28 | **36** | **1,973** | <20 |
| Experimental | RJy18 | 28 | **45** | **924** | <20 |
| Experimental | RTs18 | 28 | **21** | **606** | <20 |
| Experimental | RQk18 | 28 | **63** | **1,406** | <20 |
| Control | RFv17 | 41 | **59** | **8,302** | <20 |
| Control | RTh18 | 41 | <20 | **1,028** | <20 |
| Control | RMl18 | 41 | **32** | **1,278** | <20 |
| Control | RMu18 | 41 | **59** | **3,987** | <20 |
| Control | RYy18 | 41 | **30** | **10,256** | <20 |
| Control | RUu18 | 41 | **72** | **4,679** | <20 |
| Experimental | LJ66 | 41 | **139** | **28,893** | <20 |
| Experimental | RNp18 | 41 | <20 | **2,607** | **21** |
| Experimental | RNz18 | 41 | **21** | **5,578** | <20 |
| Experimental | RJy18 | 41 | **31** | **1,133** | <20 |
| Experimental | RTs18 | 41 | **27** | **2,559** | <20 |
| Experimental | RQk18 | 41 | **25** | **4,312** | <20 |

| | | | | | |
|---|---|---|---|---|---|
| Control | RFv17 | 43 | **32** | **4,342** | <20 |
| Control | RTh18 | 43 | **27** | **1,641** | <20 |
| Control | RMl18 | 43 | <20 | **787** | <20 |
| Control | RMu18 | 43 | **37** | **1,995** | <20 |
| Control | RYy18 | 43 | **30** | **8,230** | <20 |
| Control | RUu18 | 43 | **39** | **3,110** | <20 |
| Experimental | LJ66 | 43 | **116** | **18,270** | <20 |
| Experimental | RNp18 | 43 | <20 | **1,221** | <20 |
| Experimental | RNz18 | 43 | **21** | **9,555** | <20 |
| Experimental | RJy18 | 43 | **22** | **696** | <20 |
| Experimental | RTs18 | 43 | <20 | **771** | <20 |
| Experimental | RQk18 | 43 | **21** | **3,320** | <20 |

Values are the serum dilution when relative luminescence units (RLUs) were reduced 50% when compared against virus control wells.

**Table S5. FP Competition Neutralization Assay**

| Study Group# | Animal ID | Week | FP-10 Presence (+/-) | Virus | Dilution | % Neutralization |
|---|---|---|---|---|---|---|
| Experimental | LJ66 | 26 | - | CH505TF | 43740 | 0 |
| Experimental | LJ66 | 26 | - | CH505TF | 14580 | 0 |
| Experimental | LJ66 | 26 | - | CH505TF | 4860 | 0 |
| Experimental | LJ66 | 26 | - | CH505TF | 1620 | 0 |
| Experimental | LJ66 | 26 | - | CH505TF | 540 | 0.15709717035 |
| Experimental | LJ66 | 26 | - | CH505TF | 180 | 0.46214242643 |
| Experimental | LJ66 | 26 | - | CH505TF | 60 | 0.75534501646 |
| Control | RFv17 | 26 | - | CH505TF | 43740 | 0 |
| Control | RFv17 | 26 | - | CH505TF | 14580 | 0 |
| Control | RFv17 | 26 | - | CH505TF | 4860 | 0 |
| Control | RFv17 | 26 | - | CH505TF | 1620 | 0.21892452478 |
| Control | RFv17 | 26 | - | CH505TF | 540 | 0.51192253901 |
| Control | RFv17 | 26 | - | CH505TF | 180 | 0.79519183717 |
| Control | RFv17 | 26 | - | CH505TF | 60 | 0.94285010718 |
| Control | RMl18 | 26 | - | CH505TF | 43740 | 0 |
| Control | RMl18 | 26 | - | CH505TF | 14580 | 0 |
| Control | RMl18 | 26 | - | CH505TF | 4860 | 0 |
| Control | RMl18 | 26 | - | CH505TF | 1620 | 0.01366680031 |
| Control | RMl18 | 26 | - | CH505TF | 540 | 0.33030468535 |
| Control | RMl18 | 26 | - | CH505TF | 180 | 0.72027163120 |
| Control | RMl18 | 26 | - | CH505TF | 60 | 0.95726133611 |
| Control | RMu18 | 26 | - | CH505TF | 43740 | 0 |
| Control | RMu18 | 26 | - | CH505TF | 14580 | 0 |
| Control | RMu18 | 26 | - | CH505TF | 4860 | 0 |
| Control | RMu18 | 26 | - | CH505TF | 1620 | 0.02980555827 |
| Control | RMu18 | 26 | - | CH505TF | 540 | 0.29166258883 |
| Control | RMu18 | 26 | - | CH505TF | 180 | 0.65032943650 |
| Control | RMu18 | 26 | - | CH505TF | 60 | 0.87565832052 |
| Control | RYy18 | 26 | - | CH505TF | 43740 | 0.03707936467 |
| Control | RYy18 | 26 | - | CH505TF | 14580 | 0 |
| Control | RYy18 | 26 | - | CH505TF | 4860 | 0.00593838100 |
| Control | RYy18 | 26 | - | CH505TF | 1620 | 0.18755623466 |
| Control | RYy18 | 26 | - | CH505TF | 540 | 0.45850552323 |
| Control | RYy18 | 26 | - | CH505TF | 180 | 0.72470410698 |

| | | | | | | |
|---|---|---|---|---|---|---|
| Control | RYy18 | 26 | - | CH505TF | 60 | 0.894956638137603 |
| Control | RUu18 | 26 | - | CH505TF | 43740 | 0 |
| Control | RUu18 | 26 | - | CH505TF | 14580 | 0 |
| Control | RUu18 | 26 | - | CH505TF | 4860 | 0.07072071929 |
| Control | RUu18 | 26 | - | CH505TF | 1620 | 0.12550157378 |
| Control | RUu18 | 26 | - | CH505TF | 540 | 0.36599179802 |
| Control | RUu18 | 26 | - | CH505TF | 180 | 0.67096886217 |
| Control | RUu18 | 26 | - | CH505TF | 60 | 0.87556739794 |
| NA | VRC34.01 | NA | - | CH505TF | 0.002286 | 0 |
| NA | VRC34.01 | NA | - | CH505TF | 0.006858 | 0 |
| NA | VRC34.01 | NA | - | CH505TF | 0.020576 | 0.05799155809 |
| NA | VRC34.01 | NA | - | CH505TF | 0.061728 | 0.22483449249 |
| NA | VRC34.01 | NA | - | CH505TF | 0.185185 | 0.45373208777 |
| NA | VRC34.01 | NA | - | CH505TF | 0.555555 | 0.72631798278 |
| NA | VRC34.01 | NA | - | CH505TF | 1.666666 | 0.85426878356 |
| Experimental | LJ66 | 26 | + | CH505TF | 43740 | 0.13686689628 |
| Experimental | LJ66 | 26 | + | CH505TF | 14580 | 0 |
| Experimental | LJ66 | 26 | + | CH505TF | 4860 | 0 |
| Experimental | LJ66 | 26 | + | CH505TF | 1620 | 0 |
| Experimental | LJ66 | 26 | + | CH505TF | 540 | 0.16528020255 |
| Experimental | LJ66 | 26 | + | CH505TF | 180 | 0.50828563581 |
| Experimental | LJ66 | 26 | + | CH505TF | 60 | 0.84156235300 |
| Control | RFv17 | 26 | + | CH505TF | 43740 | 0.14323147689 |
| Control | RFv17 | 26 | + | CH505TF | 14580 | 0 |
| Control | RFv17 | 26 | + | CH505TF | 4860 | 0.01343949386 |
| Control | RFv17 | 26 | + | CH505TF | 1620 | 0.15641525099 |
| Control | RFv17 | 26 | + | CH505TF | 540 | 0.51555944221 |
| Control | RFv17 | 26 | + | CH505TF | 180 | 0.79844231940 |
| Control | RFv17 | 26 | + | CH505TF | 60 | 0.95876155869 |
| Control | RMl18 | 26 | + | CH505TF | 43740 | 0.06162846129 |
| Control | RMl18 | 26 | + | CH505TF | 14580 | 0 |
| Control | RMl18 | 26 | + | CH505TF | 4860 | 0 |
| Control | RMl18 | 26 | + | CH505TF | 1620 | 0 |
| Control | RMl18 | 26 | + | CH505TF | 540 | 0.24551937945 |
| Control | RMl18 | 26 | + | CH505TF | 180 | 0.69031264107 |
| Control | RMl18 | 26 | + | CH505TF | 60 | 0.93484892013 |
| Control | RMu18 | 26 | + | CH505TF | 43740 | 0.10754436422 |
| Control | RMu18 | 26 | + | CH505TF | 14580 | 0 |

| | | | | | | |
|---|---|---|---|---|---|---|
| Control | RMu18 | 26 | + | CH505TF | 4860 | 0 |
| Control | RMu18 | 26 | + | CH505TF | 1620 | 0.03276054212 |
| Control | RMu18 | 26 | + | CH505TF | 540 | 0.36940139477 |
| Control | RMu18 | 26 | + | CH505TF | 180 | 0.68810776851 |
| Control | RMu18 | 26 | + | CH505TF | 60 | 0.92155149280 |
| Control | RYy18 | 26 | + | CH505TF | 43740 | 0.15391488004 |
| Control | RYy18 | 26 | + | CH505TF | 14580 | 0.03594283242 |
| Control | RYy18 | 26 | + | CH505TF | 4860 | 0.10322554166 |
| Control | RYy18 | 26 | + | CH505TF | 1620 | 0.24324631495 |
| Control | RYy18 | 26 | + | CH505TF | 540 | 0.50623987775 |
| Control | RYy18 | 26 | + | CH505TF | 180 | 0.77641632439 |
| Control | RYy18 | 26 | + | CH505TF | 60 | 0.93846309269 |
| Control | RUu18 | 26 | + | CH505TF | 43740 | 0.15596063809 |
| Control | RUu18 | 26 | + | CH505TF | 14580 | 0.19096583142 |
| Control | RUu18 | 26 | + | CH505TF | 4860 | 0.24688321815 |
| Control | RUu18 | 26 | + | CH505TF | 1620 | 0.32280357249 |
| Control | RUu18 | 26 | + | CH505TF | 540 | 0.52101479701 |
| Control | RUu18 | 26 | + | CH505TF | 180 | 0.80817102810 |
| Control | RUu18 | 26 | + | CH505TF | 60 | 0.95742272531 |
| NA | VRC34.01 | NA | + | CH505TF | 0.002286 | 0.24597399235 |
| NA | VRC34.01 | NA | + | CH505TF | 0.006858 | 0.15186912199 |
| NA | VRC34.01 | NA | + | CH505TF | 0.020576 | 0.11163588032 |
| NA | VRC34.01 | NA | + | CH505TF | 0.061728 | 0.10436207391 |
| NA | VRC34.01 | NA | + | CH505TF | 0.185185 | 0.02707788087 |
| NA | VRC34.01 | NA | + | CH505TF | 0.555555 | 0.16050676710 |
| NA | VRC34.01 | NA | + | CH505TF | 1.666666 | 0.57436361591 |
| Experimental | RJy18 | 41 | - | BG505/T332N.N611A | 43740 | 0.09611230738 |
| Experimental | RJy18 | 41 | - | BG505/T332N.N611A | 14580 | 0 |
| Experimental | RJy18 | 41 | - | BG505/T332N.N611A | 4860 | 0 |
| Experimental | RJy18 | 41 | - | BG505/T332N.N611A | 1620 | 0.17822650268 |
| Experimental | RJy18 | 41 | - | BG505/T332N.N611A | 540 | 0.42299694641 |
| Experimental | RJy18 | 41 | - | BG505/T332N.N611A | 180 | 0.58746720504 |
| Experimental | RJy18 | 41 | - | BG505/T332N.N611A | 60 | 0.74093246843 |
| Experimental | RTs18 | 41 | - | BG505/T332N.N611A | 43740 | 0.02790552366 |
| Experimental | RTs18 | 41 | - | BG505/T332N.N611A | 14580 | 0 |
| Experimental | RTs18 | 41 | - | BG505/T332N.N611A | 4860 | 0.04072452911 |
| Experimental | RTs18 | 41 | - | BG505/T332N.N611A | 1620 | 0.23869350953 |
| Experimental | RTs18 | 41 | - | BG505/T332N.N611A | 540 | 0.50087845123 |
| Experimental | RTs18 | 41 | - | BG505/T332N.N611A | 180 | 0.72242956433 |

| | | | | | | |
|---|---|---|---|---|---|---|
| Experimental | RTs18 | 41 | - | BG505/T332N.N611A | 60 | 0.80901831814 |
| Experimental | RQk18 | 41 | - | BG505/T332N.N611A | 43740 | 0.39397278312 |
| Experimental | RQk18 | 41 | - | BG505/T332N.N611A | 14580 | 0.48418955734 |
| Experimental | RQk18 | 41 | - | BG505/T332N.N611A | 4860 | 0.71638286365 |
| Experimental | RQk18 | 41 | - | BG505/T332N.N611A | 1620 | 0.85203454681 |
| Experimental | RQk18 | 41 | - | BG505/T332N.N611A | 540 | 0.93240729232 |
| Experimental | RQk18 | 41 | - | BG505/T332N.N611A | 180 | 0.97955099888 |
| Experimental | RQk18 | 41 | - | BG505/T332N.N611A | 60 | 0.99319356496 |
| Control | RFv17 | 41 | - | BG505/T332N.N611A | 43740 | 0.01472371616 |
| Control | RFv17 | 41 | - | BG505/T332N.N611A | 14580 | 0 |
| Control | RFv17 | 41 | - | BG505/T332N.N611A | 4860 | 0 |
| Control | RFv17 | 41 | - | BG505/T332N.N611A | 1620 | 0 |
| Control | RFv17 | 41 | - | BG505/T332N.N611A | 540 | 0 |
| Control | RFv17 | 41 | - | BG505/T332N.N611A | 180 | 0.28343909460 |
| Control | RFv17 | 41 | - | BG505/T332N.N611A | 60 | 0.69727528948 |
| NA | VRC34.01 | NA | - | BG505/T332N.N611A | 0.002286 | 0.50075751722 |
| NA | VRC34.01 | NA | - | BG505/T332N.N611A | 0.006858 | 0.56170826012 |
| NA | VRC34.01 | NA | - | BG505/T332N.N611A | 0.020576 | 0.79189406180 |
| NA | VRC34.01 | NA | - | BG505/T332N.N611A | 0.061728 | 0.88315086854 |
| NA | VRC34.01 | NA | - | BG505/T332N.N611A | 0.185185 | 0.93056909531 |
| NA | VRC34.01 | NA | - | BG505/T332N.N611A | 0.555555 | 0.93800653715 |
| NA | VRC34.01 | NA | - | BG505/T332N.N611A | 1.666666 | 0.96547065166 |
| Experimental | RJy18 | 41 | + | BG505/T332N.N611A | 43740 | 0.30919803952 |
| Experimental | RJy18 | 41 | + | BG505/T332N.N611A | 14580 | 0.04108733115 |
| Experimental | RJy18 | 41 | + | BG505/T332N.N611A | 4860 | 0.01847267059 |
| Experimental | RJy18 | 41 | + | BG505/T332N.N611A | 1620 | 0.08873533255 |
| Experimental | RJy18 | 41 | + | BG505/T332N.N611A | 540 | 0.28573684086 |
| Experimental | RJy18 | 41 | + | BG505/T332N.N611A | 180 | 0.61661230234 |
| Experimental | RJy18 | 41 | + | BG505/T332N.N611A | 60 | 0.84615715375 |
| Experimental | RTs18 | 41 | + | BG505/T332N.N611A | 43740 | 0.30919803952 |
| Experimental | RTs18 | 41 | + | BG505/T332N.N611A | 14580 | 0.02790552366 |
| Experimental | RTs18 | 41 | + | BG505/T332N.N611A | 4860 | 0.07676286519 |
| Experimental | RTs18 | 41 | + | BG505/T332N.N611A | 1620 | 0.15077448157 |
| Experimental | RTs18 | 41 | + | BG505/T332N.N611A | 540 | 0.37534894501 |
| Experimental | RTs18 | 41 | + | BG505/T332N.N611A | 180 | 0.81656460059 |
| Experimental | RTs18 | 41 | + | BG505/T332N.N611A | 60 | 0.95829926465 |
| Experimental | RQk18 | 41 | + | BG505/T332N.N611A | 43740 | 0.70621231309 |
| Experimental | RQk18 | 41 | + | BG505/T332N.N611A | 14580 | 0.56920616897 |
| Experimental | RQk18 | 41 | + | BG505/T332N.N611A | 4860 | 0.70828028473 |

| | | | | | | |
|---|---|---|---|---|---|---|
| Experimental | RQk18 | 41 | + | BG505/T332N.N611A | 1620 | 0.84037650789 |
| Experimental | RQk18 | 41 | + | BG505/T332N.N611A | 540 | 0.94124756872 |
| Experimental | RQk18 | 41 | + | BG505/T332N.N611A | 180 | 0.98453831760 |
| Experimental | RQk18 | 41 | + | BG505/T332N.N611A | 60 | 0.99752058397 |
| Control | RFv17 | 41 | + | BG505/T332N.N611A | 43740 | 0.31536567422 |
| Control | RFv17 | 41 | + | BG505/T332N.N611A | 14580 | 0.01677959440 |
| Control | RFv17 | 41 | + | BG505/T332N.N611A | 4860 | 0 |
| Control | RFv17 | 41 | + | BG505/T332N.N611A | 1620 | 0 |
| Control | RFv17 | 41 | + | BG505/T332N.N611A | 540 | 0 |
| Control | RFv17 | 41 | + | BG505/T332N.N611A | 180 | 0.33846407084 |
| Control | RFv17 | 41 | + | BG505/T332N.N611A | 60 | 0.65482745067 |
| NA | VRC34.01 | NA | + | BG505/T332N.N611A | 0.002286 | 0.53159569071 |
| NA | VRC34.01 | NA | + | BG505/T332N.N611A | 0.006858 | 0.43908117023 |
| NA | VRC34.01 | NA | + | BG505/T332N.N611A | 0.020576 | 0.57900182408 |
| NA | VRC34.01 | NA | + | BG505/T332N.N611A | 0.061728 | 0.57972742817 |
| NA | VRC34.01 | NA | + | BG505/T332N.N611A | 0.185185 | 0.57428539755 |
| NA | VRC34.01 | NA | + | BG505/T332N.N611A | 0.555555 | 0.69079322635 |
| NA | VRC34.01 | NA | + | BG505/T332N.N611A | 1.666666 | 0.76242244266 |

### Table S6. CryoEM data collection, processing and model building statistics.

| **Map** | **Autologous Boost RUu18 wk 14 Polyclonal Fab FP1** | **Autologous Boost RUu18 wk 14 Polyclonal Fab V1V2V3-1** | **Autologous Boost RUu18 wk 14 Polyclonal Fab V1V2V3-2** | **Autologous Boost RUu18 wk 14 Polyclonal Fab V1V2V3-5** |
|---|---|---|---|---|
| EMDB | EMD-49415 | EMD-49416 | EMD-49417 | EMD-49418 |
| **Data collection** | | | | |
| Microscope | TFS Talos Arctica | TFS Talos Arctica | TFS Talos Arctica | TFS Talos Arctica |
| Voltage (kV) | 200 | 200 | 200 | 200 |
| Detector | Gatan K2 Summit | Gatan K2 Summit | Gatan K2 Summit | Gatan K2 Summit |
| Recording mode | Counting | Counting | Counting | Counting |
| Nominal magnification | 36,000x | 36,000x | 36,000x | 36,000x |
| Movie micrograph pixelsize (Å) | 1.15 | 1.15 | 1.15 | 1.15 |
| Total dose ($e^-/Å^2$) | 49.92 | 49.92 | 49.92 | 49.92 |
| Defocus range (µm) | -1.8 to -0.8 | -1.8 to -0.8 | -1.8 to -0.8 | -1.8 to -0.8 |
| **EM data processing** | | | | |
| Number of movie micrographs | 7500 | 7500 | 7500 | 7500 |
| Number of molecular projection images in map | 143,186 | 47,505 | 87,728 | 171,960 |
| Symmetry | C1 | C1 | C1 | C1 |
| Map pixel size | 1.15 | 1.15 | 1.15 | 1.15 |
| Map resolution (FSC 0.143; Å) | 3.7 | 4.0 | 3.9 | 3.8 |
| Map sharpening B-factor ($Å^2$) | -120.2 | -112.7 | -128.5 | -125.5 |
| **Structure building and validation** | | | | |
| *Model Composition* | | | | |
| Non-hydrogen atoms | 15017 | 14897 | 15064 | 14784 |
| Protein residues | 1873 | 1856 | 1880 | 1851 |
| ligands | 64 | 67 | 65 | 60 |
| MolProbity score | 1.00 | 1.05 | 1.13 | 0.92 |
| Clashscore | 1.05 | 1.09 | 1.28 | 0.69 |
| EMRinger score | 2.88 | 2.21 | 2.02 | 2.10 |
| d FSC model (0.5; Å) | 3.9 | 4.2 | 4.1 | 4.0 |
| *RMSD from ideal* | | | | |
| Bond length (Å) | 0.021 | 0.021 | 0.021 | 0.021 |
| Bond angles (˚) | 1.712 | 1.707 | 1.722 | 1.717 |
| Rama Outliers (%) | 0.00 | 0.00 | 0.00 | 0.00 |
| Side chain rotamer outliers (%) | 0.00 | 0.00 | 0.00 | 0.27 |
| Cβ outliers (%) | 0.00 | 0.00 | 0.00 | 0.00 |
| PDB | 9NHL | 9NHM | 9NHN | 9NHO |

### Table S6 continued

| Map | Autologous Boost RUu18 wk 14 Polyclonal Fab Base 1 | Autologous Boost RUu18 wk 14 Polyclonal Fab Base 4 | Heterologous Boost RQk18 wk 43 Polyclonal Fab Base A | Heterologous Boost RQk18 wk 43 Polyclonal Fab Base C |
|---|---|---|---|---|
| EMDB | EMD-49457 | EMD-49414 | EMD-49411 | EMD-49412 |
| **Data collection** | | | | |
| Microscope | TFS Talos Arctica | TFS Talos Arctica | TFS Titan Krios | TFS Titan Krios |
| Voltage (kV) | 200 | 200 | 300 | 300 |
| Detector | Gatan K2 Summit | Gatan K2 Summit | Gatan K2 Summit | Gatan K2 Summit |
| Recording mode | Counting | Counting | Counting | Counting |
| Nominal magnification | 36,000x | 36,000x | 130,000x | 130,000x |
| Movie micrograph pixelsize (Å) | 1.15 | 1.15 | 1.045 | 1.045 |
| Total dose ($e^-$/Å$^2$) | 49.92 | 49.92 | 50.29 | 50.29 |
| Defocus range (µm) | -1.8 to -0.8 | -1.8 to -0.8 | -1.8 to -0.8 | -1.8 to -0.8 |
| **EM data processing** | | | | |
| Number of movie micrographs | 7500 | 7500 | 11745 | 11745 |
| Number of molecular projection images in map | 181,253 | 110,868 | 357902 | 145,407 |
| Symmetry | C1 | C1 | C1 | C1 |
| Map pixel size | 1.15 | 1.15 | 1.045 | 1.045 |
| Map resolution (FSC 0.143; Å) | 3.8 | 3.89 | 3.0 | 3.1 |
| Map sharpening B-factor (Å$^2$) | -124.0 | -116.9 | -94.97 | -89.0 |
| **Structure building and validation** | | | | |
| *Number of atoms in deposited model* | | | | |
| Non-hydrogen atoms | 14956 | 14953 | 15606 | 15028 |
| Protein residues | 1879 | 1859 | 1889 | 1844 |
| ligands | 58 | 69 | 97 | 75 |
| MolProbity score | 1.11 | 1.07 | 0.59 | 0.90 |
| Clashscore | 1.39 | 0.88 | 0.23 | 0.88 |
| EMRinger score | 2.61 | 2.42 | 4.40 | 4.05 |
| d FSC model (0.5; Å) | 3.9 | 4.0 | 3.1 | 3.2 |
| *RMSD from ideal* | | | | |
| Bond length (Å) | 0.021 | 0.021 | 0.005 | 0.005 |
| Bond angles (˚) | 1.706 | 1.731 | 0.935 | 0.941 |
| Rama Outliers (%) | 0.00 | 0.00 | 0.00 | 0.00 |
| Side chain rotamer outliers (%) | 0.00 | 0.14 | 0.00 | 0.00 |
| Cβ outliers (%) | 0.00 | 0.00 | 0.00 | 0.00 |
| PDB | 9NI9 | 9NHK | 9NHH | 9NHI |

### Table S6 continued

| Map | Heterologous Boost RQk18 wk 43 Polyclonal Fab FP-A | Heterologous Boost RQk_FP_mAb_05 |
|---|---|---|
| EMDB | EMD-49413 | |
| **Data collection** | | |
| Microscope | TFS Titan Krios | TFS Glacios |
| Voltage (kV) | 300 | 200 |
| Detector | Gatan K2 Summit | TFS Falcon 4 |
| Recording mode | Counting | Counting |
| Nominal magnification | 130,000x | 190,000x |
| Movie micrograph pixelsize (Å) | 1.045 | 0.725 |
| Total dose ($e^-$/Å$^2$) | 50.29 | 45 |
| Defocus range (µm) | -1.8 to -0.8 | -2.0 to -1.0 |
| **EM data processing** | | |
| Number of movie micrographs | 11,745 | 16,477 |
| Number of molecular projection images in map | 254,630 | 111,683 |
| Symmetry | C1 | C1 |
| Map pixel size | 1.045 | 0.725 |
| Map resolution (FSC 0.143; Å) | 3.0 | 3.9 |
| Map sharpening B-factor (Å$^2$) | -90.3 | -93.5 |
| **Structure building and validation** | | |
| *Number of atoms in deposited model* | | *NA* |
| Non-hydrogen atoms | 15437 | |
| Protein residues | 1905 | |
| ligands | 76 | |
| MolProbity score | 0.94 | |
| Clashscore | 0.85 | |
| EMRinger score | 3.42 | |
| d FSC model (0.5; Å) | 3.2 | |
| *RMSD from ideal* | | |
| Bond length (Å) | 0.005 | |
| Bond angles (˚) | 0.973 | |
| Rama Outliers (%) | 0.00 | |
| Side chain rotamer outliers (%) | 0.00 | |
| Cβ outliers (%) | 0.00 | |
| PDB | 9NHJ | |

**Table S7. RQk_FP_mAb Design – Heavy chain**

| H-chain name | Additional mutations | Top Score Hit #ID | Assigned FWRH1 region | Assigned JH region |
|---|---|---|---|---|
| RQk_FP1_HC_v1 | | 980571, 979518 | IGHV4-117*01 | IGHJ4-3*01 |
| RQk_FP1_HC_v2 | G59R | 980571, 979518 | IGHV4-117*01 | IGHJ4-3*01 |
| RQk_FP1_HC_v3 | | 980571, 979518 | IGHV4-117*01 | IGHJ4-3*01 |
| RQk_FP1_HC_v4 | D114R | 980571, 979518 | IGHV4-117*01 | IGHJ4-3*01 |

**Table S8. RQk_FP_mAb Design – Light chain**

| L-chain name | Additional mutations | Top Score Hit #ID |
|---|---|---|
| RQk_FP1_LC_v1 | | flexible CDRL3, ordered (SSSPV) |
| RQk_FP1_LC_v2 | S108N, S114N, V116P | Based on top pick CDRH3s (NSNPP) |

**Table S9. RQk_FP_mAb Design – Heavy chain and Light Chain Pairing**

| # Antibody | H-chain | L-chain |
|---|---|---|
| RQk_FP1_Ab1_IgG | RQk_FP1_HC_v1 | RQk_FP1_LC_v1 |
| RQk_FP1_Ab2_IgG | RQk_FP1_HC_v2 | RQk_FP1_LC_v1 |
| RQk_FP1_Ab3_IgG | RQk_FP1_HC_v3 | RQk_FP1_LC_v1 |
| RQk_FP1_Ab4_IgG | RQk_FP1_HC_v4 | RQk_FP1_LC_v1 |
| RQk_FP1_Ab5_IgG | RQk_FP1_HC_v1 | RQk_FP1_LC_v2 |
| RQk_FP1_Ab6_IgG | RQk_FP1_HC_v2 | RQk_FP1_LC_v2 |
| RQk_FP1_Ab7_IgG | RQk_FP1_HC_v3 | RQk_FP1_LC_v2 |
| RQk_FP1_Ab8_IgG | RQk_FP1_HC_v4 | RQk_FP1_LC_v2 |

**Table S10. AMC016 and BG505 pseudovirus neutralization data for RQk_FP_mAbs**

| mAb | SVA-MLV | BG505 | BG505/ T332N | BG505/ T332N. N611A | BG505/ T332N. T465N | BG505/ T332N. 133aN+ 136aA | BG505/ T332N. S241N. P291T | AMC016 |
|---|---|---|---|---|---|---|---|---|
| RQk_FP_mAb _01 | >50.0 | >50.0 | >50.0 | >50.0 | >50.0 | >50.0 | >50.0 | >50.0 |
| RQk_FP_mAb _03 | >50.0 | >50.0 | >50.0 | >50.0 | >50.0 | >50.0 | >50.0 | >50.0 |
| RQk_FP_mAb _05 | >50.0 | >50.0 | >50.0 | >50.0 | >50.0 | >50.0 | >50.0 | >50.0 |
| RQk_FP_mAb _07 | >50.0 | >50.0 | >50.0 | >50.0 | >50.0 | >50.0 | >50.0 | >50.0 |

**Table S11. FP-sensitive pseudovirus neutralization data for RQk_FP_mAbs**

| mAb | 3988.25 | 0077.v1.c16 | 25710-2.43 | 286_36 | BL01.DG | CNE19 | CNE56 | KER2008.vrc12 | Q23.17 |
|---|---|---|---|---|---|---|---|---|---|
| RQk_FP_mAb_01 | >50.0 | >50.0 | >50.0 | >50.0 | >50.0 | >50.0 | >50.0 | >50.0 | >50.0 |
| RQk_FP_mAb_03 | >50.0 | >50.0 | >50.0 | >50.0 | >50.0 | >50.0 | >50.0 | >50.0 | >50.0 |
| RQk_FP_mAb_05 | >50.0 | >50.0 | >50.0 | >50.0 | >50.0 | >50.0 | >50.0 | >50.0 | >50.0 |
| RQk_FP_mAb_07 | >50.0 | >50.0 | >50.0 | >50.0 | >50.0 | >50.0 | >50.0 | >50.0 | >50.0 |

**Table S12: FNA Phenotyping Staining Panel**

| | |
|---|---|
| Viability | e506 |
| CD8a | BV510 |
| PD-1 (Clone EH12.2H7) | BV605 |
| CD4 (Clone OKT-4) | BV785 |
| CD20 (Clone 2H7) | Ax488 |
| IgG (Clone G18-145) | AF700 |
| CXCR5 (Clone Mu5UBEE) | PECy7 |
| IgM (Clone G20-127) | PerCP-Cy5.5 |
| CD71 (Clone L01.1) | PE-CF594 |
| CD38 (Clone OKT) | PE |

**Table S13: 10X B cell Staining Panel**

| | |
|---|---|
| Live/Dead | APCe780 |
| CD8a | APCe780 |
| CD4 (Clone SK3) | APCe780 |
| CD16 (ebioCD16) | APCe780 |
| CD20 (Clone 2H7) | Ax488 |
| IgG (Clone G18-145) | PECy7 |
| IgM (Clone G20-127) | PerCP-Cy5.5 |
| CD71 (Clone L01.1) | PE-CF594 |
| CD38 (Clone OKT) | PE |

**Supplementary Data 01**
Bulk B cell repertoire heavy chain sequencing analysis of RQk18 PBMC from week 42

**Supplementary Data 02**
Bulk B cell repertoire light chain sequencing analysis of RQk18 PBMC from week 42

**Supplementary Data 03**
10X Genomics meta-analysis of RQk18 week 14 PBMC clonotypes.
